# Glutathione-mediated plant response to high-temperature

**DOI:** 10.1101/2022.03.28.485658

**Authors:** Avilien Dard, Alizée Weiss, Laetitia Bariat, Nathalie Picault, Frédéric Pontvianne, Christophe Riondet, Jean-Philippe Reichheld

## Abstract

Climate change induce global warming and intense heat waves that affect plant development and productivity. Among the molecular perturbations that high temperature induces in living cells is the accumulation of reactive oxygen species (ROS), which can damage macromolecules of the cell and perturb the cellular redox state. To cope with deleterious effects of ROS, plant, as other organisms, have developed strategies to scavenge ROS and to regulate their redox state. Among those, glutathione plays a major role in maintaining the cellular redox state and the function of key antioxidant enzymes like peroxidases. Here, we investigated the contribution of the redox systems in plant adaptation to high temperature. We studied two different high temperature regimes: a rise of ambient temperature to 27°C inducing a plant developmental adaptation program called thermomorphogenesis, and a 37°C treatment mimicking intense heat wave and affecting plant viability. Using the genetically encoded redox marker roGFP, we show that high temperature regimes lead to cytoplasm and nuclear oxidation and impact profoundly the glutathione pool rather than the glutathione redox state. Moreover, plant can restore the pool within a few hours, which likely contribute to plant adaptation to high temperature. However, conditional glutathione deficient mutants fail to adapt to intense heat waves or to induce thermomorphogenesis, suggesting that glutathione is involved in both heat adaptation mechanisms. We also evaluate by RNAseq analyses, how plant change its genome expression signature upon heat stress and identified a marked genome expression deviation in mutant deficient in glutathione antioxidant which might contribute to its sensitivity to high temperature. Thus, we define glutathione as a major antioxidant molecule acting in the adaptation of plant to rise of temperature.

## INTRODUCTION

Climate change is impacting all levels of the ecological pyramid. At the base of the pyramid, plants, as sessile organisms are particularly prone to the multifactorial environmental stress induced by the rise of global temperature. Among stress induced by temperature rise account systemic water deficit, increased pest ravage, or increased salt concentrations in flooding regions. Moreover, climate change induces more frequent and intense heat waves which impact plant survival (Liu et al., 2016). The injuries all these stresses induce in plant cell are diverse and act at all plant organs and cell compartments. A common feature of plant subjected to environmental stress is the generation of reactive oxygen species (ROS) which are by-products of major cellular metabolic activities like plastidial, and mitochondrial electrons transfer chains, but also accumulate at excessive level upon stress conditions (Choudhury et al., 2017; Noctor, 2017). The reactivity of these species means that their accumulation must be controlled, but also that they are excellent signaling molecules that underpin plant development and responses to the environment (Considine and Foyer, 2014; Dietz et al., 2016; Waszczak et al., 2018; Mhamdi and Breusegem, 2018; Huang et al., 2019). Upon fluctuating temperatures, ROS also accumulate at the plasma membrane/apoplast interface by NADPH oxidases, superoxide dismutases and peroxidases, and can be translocated to the cytosolic compartment by aquaporins, were they likely take part to stress signaling pathways (Mittler et al., 2012; Rodrigues et al., 2017). Moreover, receptor kinase has been recently shown to sense H_2_O_2_ at the plasma membrane and to activate signaling cascades (Wu et al., 2020). Further emphasizing their role as signaling molecules, ROS and particularly H_2_O_2_, are also major regulators of gene expression, as evidenced by extensive genome expression reprogramming under conditions promoting H_2_O_2_ accumulation (Queval et al., 2007; Willems et al., 2016). While the transcriptional mechanisms controlling gene expression by ROS are not fully understood, they partially rely on nuclear ROS translocation from the cytosol and organelles, activation and redox modifications of transcription factors and chromatin remodeling proteins (Dietz, 2014; Exposito-Rodriguez et al., 2017; Martins et al., 2018).

To control ROS accumulation particularly under stress conditions, plants have developed large panels of antioxidant molecules and enzymes (Noctor, 2017). Among those molecules, ascorbate and glutathione are key players because they serve as reducing power for specific ROS detoxification enzyme like peroxidases. Moreover, their oxidized forms are regenerated by specific reduction systems (dehydroascorbate or glutathione reductase), enabling them to act as effective regulator of the cellular redox state (Foyer and Noctor, 2011). For example, the pool of cytosolic glutathione is known to be highly reduced under standard growth conditions, but is susceptible to oxidation upon stress conditions, impacting different aspects of cell metabolism (Queval et al., 2011; Babbar et al., 2020). Consistently, mutants with affected levels of glutathione or perturbed glutathione reductase activities are more sensitive (a)biotic stress conditions and are affected in different developmental processes (Cobbett et al., 1998; Vernoux et al., 2000; Ball et al., 2004; Parisy et al., 2007; Reichheld et al., 2007; Marty et al., 2009; Bashandy et al., 2010; Mhamdi et al., 2010; Shanmugam et al., 2012; Yu et al., 2013). Therefore, monitoring the glutathione redox state under stress conditions can serve as a proxy to evaluate the impact of stress on plant fitness (Meyer et al., 2007).

Plants as other organisms are equipped with sophisticated molecular mechanisms to perceive rising temperature and to signal these modifications to appropriate stress and developmental responses. The nature of these responses is intimately dependent on the intensity of the temperature regime. For example, a moderate raise of ambient temperature (e.g. 20 to 27°C in the present analyses in Arabidopsis) leads to developmental modifications like elongation of the hypocotyl, early flowering or modifications of stomata development, a process generically called thermomorphogenesis (Casal and Balasubramanian, 2019). The molecular mechanisms underlying these developmental modifications have been largely investigated. They include a network of transcriptional and posttranscriptional regulators initiated by the photoreceptor phytochrome B (PHYB) (Jung et al., 2016; Legris et al., 2016). PHYB perceives high ambient temperature by switching from an active to an inactive form, preventing degradation of transcription factors like PHYTOCHROME INTERACTING FACTORS (PIF), which play central role in inducing developmental promoting factors like auxin metabolism, flower, or stomatal development genes (Gangappa and Kumar, 2017). Such developmental adaptation does not occur under more intense high temperature regimes (e.g. 37°C in our present experiments). Among all the rescue mechanisms dedicated to the protection of plant macromolecules from heat injuries, major actors are Heat Stress Protein (HSP)-like chaperones which protein proteins from heat-induced protein aggregation (Jacob et al., 2017). Consistent with ROS accumulation under heat stress, many antioxidant proteins like peroxidases are also taking part in the protection strategies against heat stress (Volkov et al., 2006; Mittler et al., 2012; Babbar et al., 2020). The capacity of all these mechanisms to rapidly respond to raised temperature and to maintain the intracellular redox status is key for plant survival to these changing environments.

In this work, we investigate the impact of two distinct high temperature regimes on cellular redox state. Using the ratiometric redox sensitive GFP (roGFP) construct, we show that both moderate raise of ambient temperature (27-29°C) and more intense high stress (37°C) lead to oxidation of the roGFP both in the cytosol and the nuclear compartment, but with different kinetics depending on the heat treatment. Further measurement of glutathione levels showed that roGFP oxidation is mostly due to a decrease of the extractible glutathione rather than by glutathione oxidation and that plant can restore a steady state glutathione level after long term high temperature. We further demonstrate that low glutathione mutants, unable to maintain a wild-type glutathione levels fail to survive to intense heat stress conditions, indicating that glutathione level readjustment is required for plant survival to heat stress. Moreover, low glutathione mutants also fail to induce thermomorphogenesis upon moderate raise of ambient temperature, underpinning glutathione role in this development pathway. Finally, we delineate gene expression signatures induced by high temperature regimes in both wild-type and low glutathione mutants. The role of glutathione as an actor of genome expression adjustment upon heat stress is discussed.

## RESULTS

### High temperature regimes induce cytosol and nuclear oxidation

To explore the impact of heat stress on the cytosolic redox state in cotyledons and root tip, we used as a proxy wild-type plants expressing the GRX1-roGFP2 in the cytosol (Gutscher et al., 2008). Calibration was made against 100 mM H_2_O_2_ and 10 mM DTT to fully oxidize or reduce the sensor, respectively. At 20°C, the fluorescence ratio was close to the ratio measured after DTT treatment, suggesting that under unstressed conditions the redox sensor is almost fully reduced. When the temperature is shifted 27°C, the 405/488 nm ratio does not change significantly in the root, while a mild increase is observed after 45 minutes of treatment (Figure 1A, C, E and G). However, a stronger heat treatment at 37°C triggers a fast oxidation of the roGFP in cotyledons, reaching full oxidation after 30 minutes treatments (Figure 1B and F). Increased ratio was also observed in the root tip on the same treatments but at a lesser extent (Figure 1D and H). Calibration against 100 mM H_2_O_2_ to fully oxidize and 10 mM DTT to fully reduce the sensor (Figure 1A and B) and an assumed cytosolic pH of 7.4 led to an estimation of the glutathione redox potential (E_GSH_) of - 309mV +/-10 at 20°C in palissadic cotyledon cells and to a less negative E_GSH_ to −247 mV +/-21 after 1 hour treatment at 37°C. Therefore, we concluded that a moderate increase of temperature (20°C to 27°C) has only a mild effect on the cytosolic redox state, but that an intense temperature increase (20°C to 37°C) impact the cytosolic redox state.

**Figure 1:**
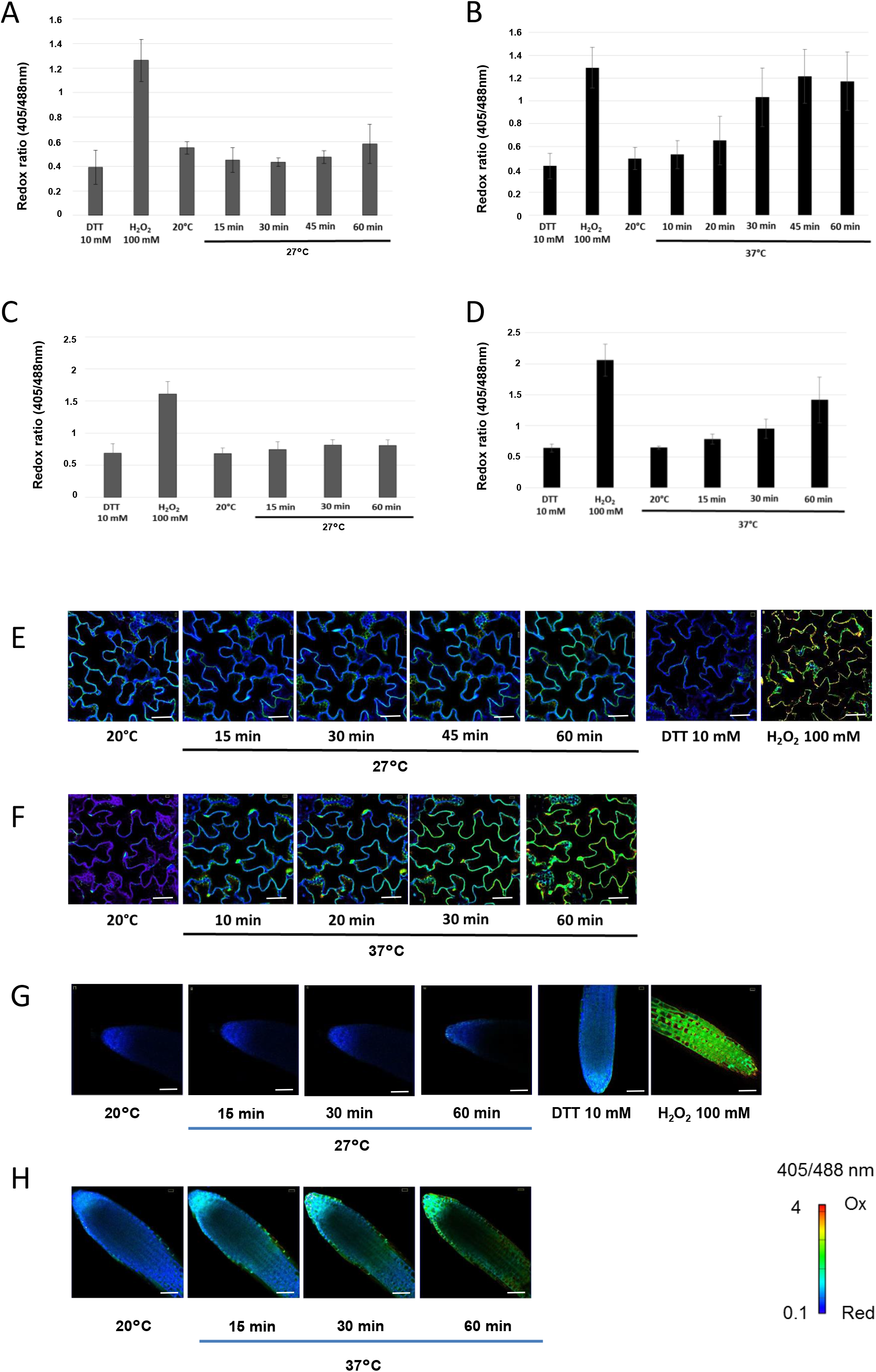
*In vivo* monitoring of the glutathione redox state upon heat stress. A-D, Confocal images of cotyledon (A-B) or root apex (C-D) cells of ten-day old Arabidopsis wild-type seedlings stably expressing the Grx1-roGFP2 construct and subjected to 27°C (A and C) or 37°C (B and D). roGFP2 fluorescence was collected at 505-530 nm after excitation with either 405 nm or 488 nm. Ratio images were calculated as the 405/488 nm fluorescence. To fully reduced or oxidize the sensor, seedlings were immersed in 10 mM DTT or 100 mM H_2_O_2_, respectively. Control samples were immersed in MS/2 liquid medium and observed at 20°C. Then, the temperature of the thermostatic chamber was increased and the roGFP2 fluorescence were monitored 10-15 min for 1-2 hours. False colors indicate the fluorescence ratio on a scale from blue (reduced) to red (oxidized). Scale bars = 50 μm. E-H, Fluorescence ratio calculated from ratio images. E-F, cotyledons at 27°C (E) and 37°C (F). G-H, root apex at 27°C (G) and 37°C (H). n=6.

The GRX1-roGFP2 constructs show localization in the nucleus because the fusion protein can diffuse through the nuclear pores (Meyer et al., 2007). This enables to study the effect of high temperature on redox state of glutathione also in the nuclear compartments (Babbar et al., 2020). When measuring the roGFP2 redox ratio by focusing our analysis on nuclei (Supplemental Figure 1), we observed a progressive oxidation upon heat stress occurring after 10 minutes, which is noticeably faster as in the cytosol (Supplemental Figure 1). Therefore, we conclude that high temperature regime has a marked effect on the glutathione redox state both in the cytosol and the nucleus, which is slightly faster in the nucleus.

### Long-term effect of high temperatures on glutathione redox state

To study the impact of high temperature regimes on the glutathione redox state at a longer term, we quantified the pools of glutathione in plants subjected to high temperature for 3 days (Figure 2). Glutathione levels were monitored in 4-day-old plants subjected to the same high temperature regimes of 27°C or 37°C. Both regimes led to a marked decrease (- 35% and −50% at 27°C and 37°C, respectively) of the total extractible glutathione within the six first hours of the treatment. However, the decrease is more pronounced at 37°C than 27°C, dropping within one while progressively decreasing after 2 hours at 27°C (Figure 2). The oxidized glutathione level (GSSG) does not accumulate at a high level during this period, and therefore the GSH/(GSH+GSSG) is not profoundly perturbed. The most pronounced change in the GSH/(GSH+GSSG) ratio is observed after 1 hour treatment at 37°C (from 87% to 73%) (Figure 2), partially reflecting the observed roGFP oxidation (Figure 1B). Interestingly, at both temperatures, the glutathione levels increased after 24 hours of treatment, to reach about the same level as untreated plants after 3 days. Therefore, we conclude that both high temperature regimes affect profoundly the reduced glutathione level rather than the oxidized glutathione and that wild-type plants are adapting their glutathione metabolism in response to high temperatures (Figure 2).

**Figure 2:**
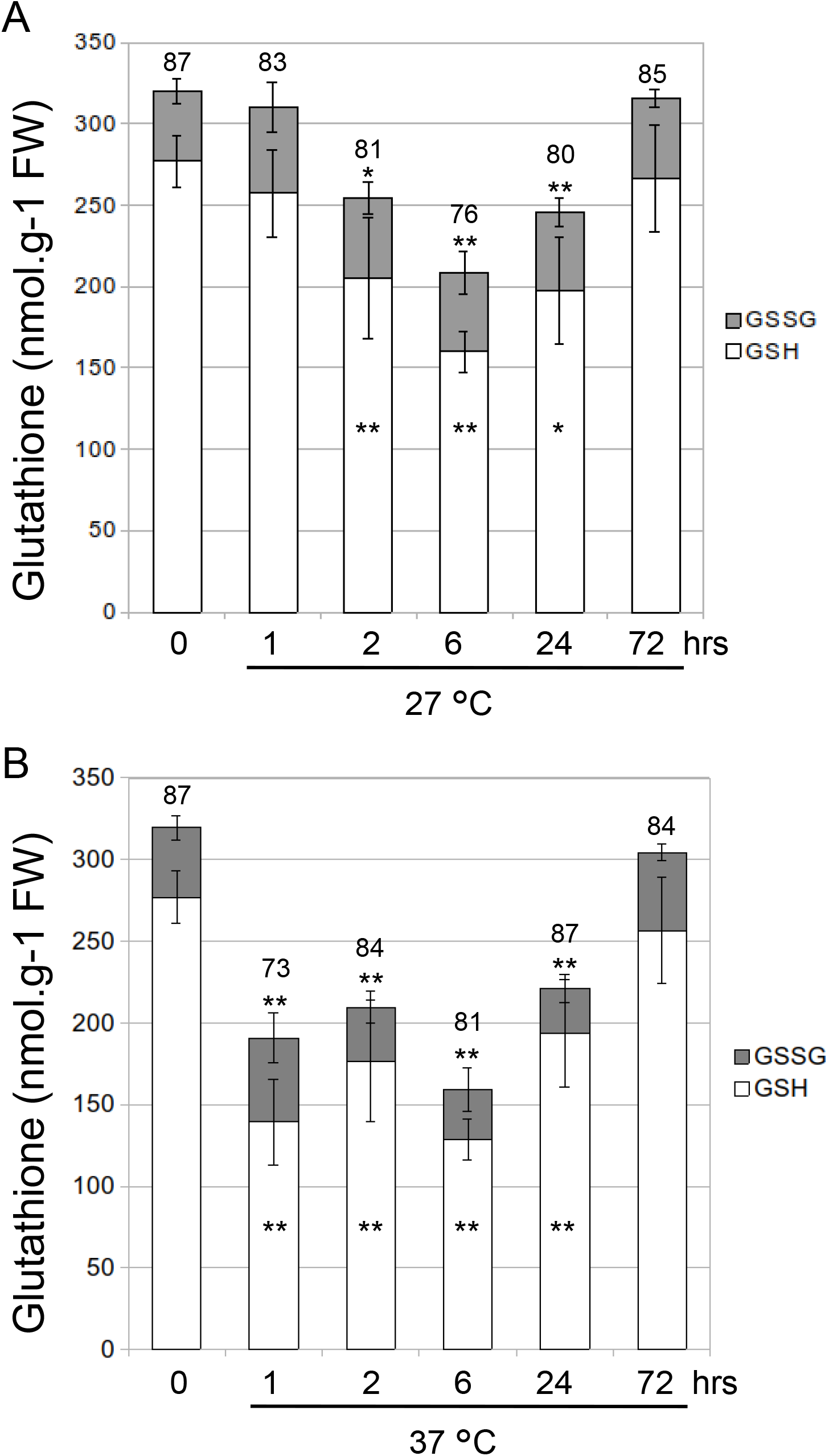
Measurement of glutathione levels upon heat stress. Total extractible glutathione was measured in ten-day old Arabidopsis wild-type seedlings subjected to 27°C (A) or 37°C (B). Reduced and oxidized glutathione levels are indicated by white and gray bars, respectively. The percentage reduction state of glutathione is indicated above bars. Asterisks indicate a significant difference (* P ≤ 0.05, ** P ≤ 0.01) between reduced glutathione levels between untreated plants (T0) and high temperature treated plants. No significant difference was observed on oxidized glutathione levels. Error bars represent SD (n = 3).

### High temperature regime changes the O_2_^.-^/H_2_O_2_ balance in the root

To evaluate the impact of high temperatures on ROS accumulation *in planta*, plants were stained with diaminobenzidine (DAB) and nitrobluetetrazolium (NBT) which react with O_2_^.-^ and H_2_O_2_ respectively. At 20°C, the highest DAB and NBT staining were found in the root (Supplemental Figure 2A and C). Staining in shoot was much lower and only restricted to leaf tip for NBT (Supplemental Figure 2C). No significant change was observed after 2 hours treatment at 37°C in the shoot, suggesting that high temperature does not generate massive O_2_^.-^ and H_2_O_2_ under these conditions. This contrasts with ROS accumulation observed by (Babbar et al., 2020), in plants subjected to a 1-hour 42°C treatment. However, in root tip, a marked change in the DAB and NBT staining was observed (Supplemental Figure 2 E-G). At 20°C, the higher DAB staining is found in the elongation zone, as NBT signal is higher in the root tip, as previously shown (Dunand et al., 2007; Tsukagoshi et al., 2010). However, at 37°C, high DAB staining is found in the meristem, while NBT is lower, suggesting that high temperatures lead to a change in O_2_^.-^/H_2_O_2_ balance (Supplemental Figure 2 E-H).

### Low glutathione impairs thermotolerance

As the steady-state level of extractible reduced glutathione is mostly affected under high temperature, we studied the response to high temperatures of the *cadmiumsensitive2* (*cad2*) and phytoalexin-deficient mutant (*pad2-1*) mutants, which contain low level of glutathione due to impaired glutathione synthesis capacities (Howden et al., 1995; Cobbett et al., 1998; Parisy et al., 2007). First, we measured the glutathione levels in the *cad2* mutant subjected to the same high temperature regimes as previously (Figure 3A). In agreement with previous reports, the total extractible glutathione level is about 12% compared to wild-type plants. Total glutathione levels decrease after 6 hours of high temperature (both 27°C and 37°C) treatments, as observed in wildtype plants (Figure 2). After 72 hours treatments, the glutathione level slightly increases at 27°C, but dramatically dropped at 37°C (Figure 3A).

**Figure 3:**
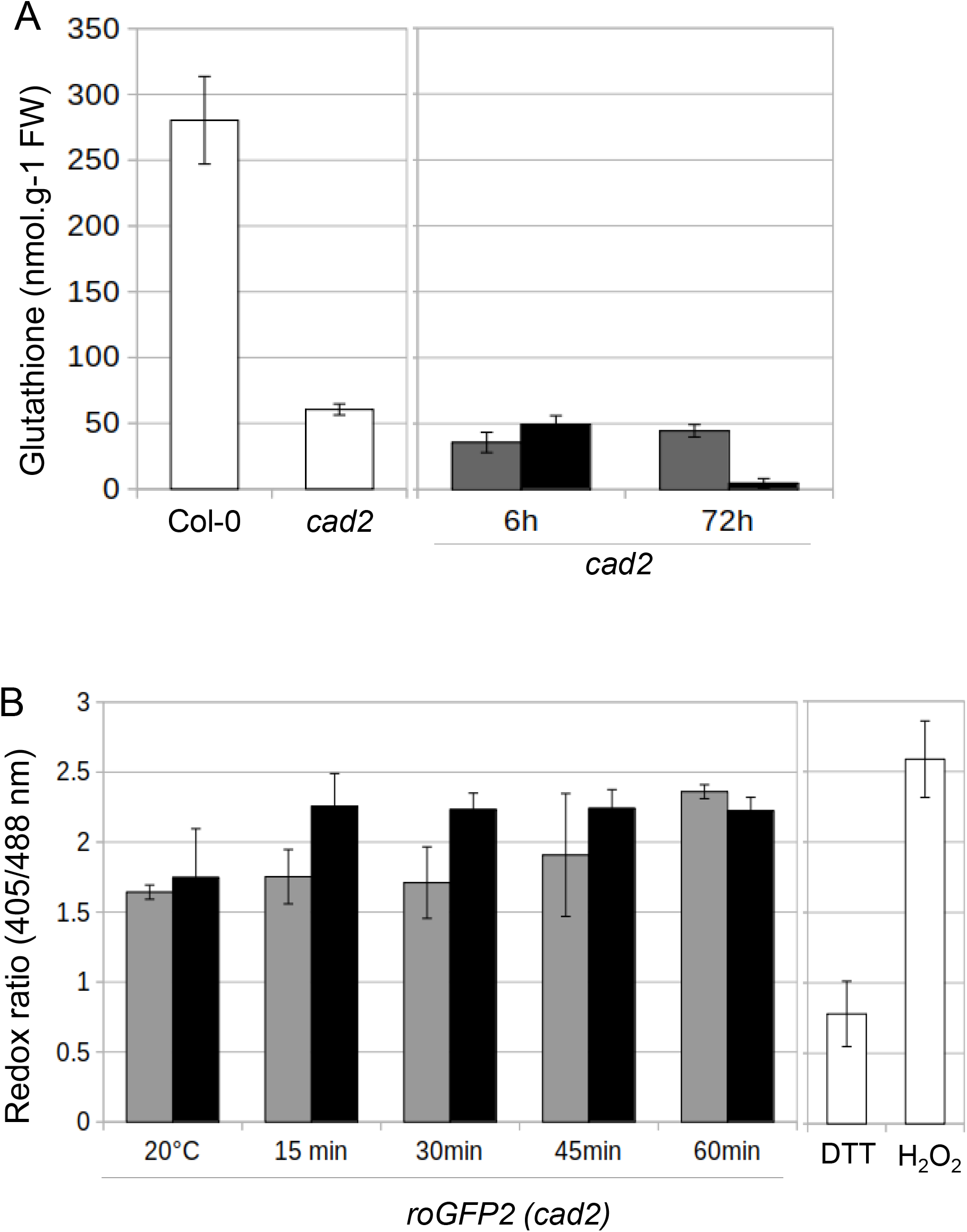
Glutathione level and redox state in low glutathione mutants. (A) Total extractible glutathione was measured in ten-day old Arabidopsis wild-type (Col-0) and *cad2* seedlings at 20°C (white bars) or subjected to 27°C (grey bars) or 37°C (black bars) for indicated times. Error bars represent SD (n = 3). (B) roGFP2 fluorescence ratio calculated from confocal images of cotyledon cells of ten-day old Arabidopsis *cad2* seedlings stably expressing the Grx1-roGFP2 construct and subjected to 27°C (grey bars) or 37°C (black bars). roGFP2 fluorescence was collected at 505-530 nm after excitation with either 405 nm or 488 nm. Ratio images were calculated as the 405/488 nm fluorescence. To fully reduced or oxidize the sensor, seedlings were immersed in 10 mM DTT or 100 mM H_2_O_2_, respectively (white bars). n=6.

We also measured the glutathione redox state in the *cad2* mutant, using the Grx1-roGFP2 construct introgressed in *cad2* (Meyer et al., 2007). As previously described, the roGFP probe is already partially oxidized under standard temperature conditions, which is due to a lower glutathione redox potential (Figure 3, Meyer et al., 2007). As expected, the roGFP is fully oxidized in the mutant after heat stress treatments (Figure 3B).

Next, we studied the response of the glutathione deficient mutants to the same high temperature regimes. A 37°C heat regime mimicking a heat wave is commonly called Thermotolerance to Moderate High Temperature (TMHT) (Yeh et al., 2012; Martins et al., 2020b). Here, seeds are germinated at standard temperature for 4 days before to be transferred to 37°C for 5 further days. Plants are switched again to 20°C to let them recover from the stress. After 11 days of recovery, the viability of the plants was assessed by the recovery of shoot growth (Figure 4A). About 80% of wild-type plants survived to the TMHT treatment while the survival rate in *cad2* and *pad2* plants were profoundly affected (Figure 4B-D). To confirm that impaired glutathione level was responsible for the loss of plant survival rate under TMHT, we treated wild-type plant with increasing concentrations of the glutathione biosynthesis inhibitor buthionine sulfoximide (BSO). This led to a progressive inhibition of plant survival rate, reaching an almost null value at 0.5 mM BSO (Figure 4E). To study whether a perturbed glutathione redox state impacts the response to TMHT, we subjected the cytosolic glutathione reductase mutant *gr1* to the same treatment (Marty et al., 2009). In contrast to glutathione deficient mutants, the survival rate of *gr1* was not perturbed (Figure 4C), indicating that the glutathione level rather than the glutathione redox state impairs plant survival to TMHT. To evaluate whether mutants deficient in different glutathione-dependent enzymes are impacted by TMHT, we subjected cytosolic glutaredoxin mutants to TMHT (Figure 4C). None of the studied *grx* mutants were affected, in contrast to the previously described *grxS17* knock-out mutants (Martins et al., 2020). As heat stress induces ROS accumulation, we also studied the behavior of mutants affected in ROS metabolism. The catalase2 (*cat2*) mutant inactivated in a major H_2_O_2_ catabolic enzyme was also affected by TMHT (Figure 4C). We also monitored the thermotolerance of GSNO and NO accumulating mutants to TMHT. The GSNO reductase mutant *hot5* was previously shown to be intolerant to a 45°C heat treatment (Lee et al., 2008). Here, we found that the *hot5-1* mutant is also intolerant to a 37°C TMHT treatment (Figure 4C). Finally, we also found the NO accumulating mutant *nox1* to be highly sensitive to TMHT, as previously described at 45°C (Lee et al., 2008) (Figure 4C).

**Figure 4:**
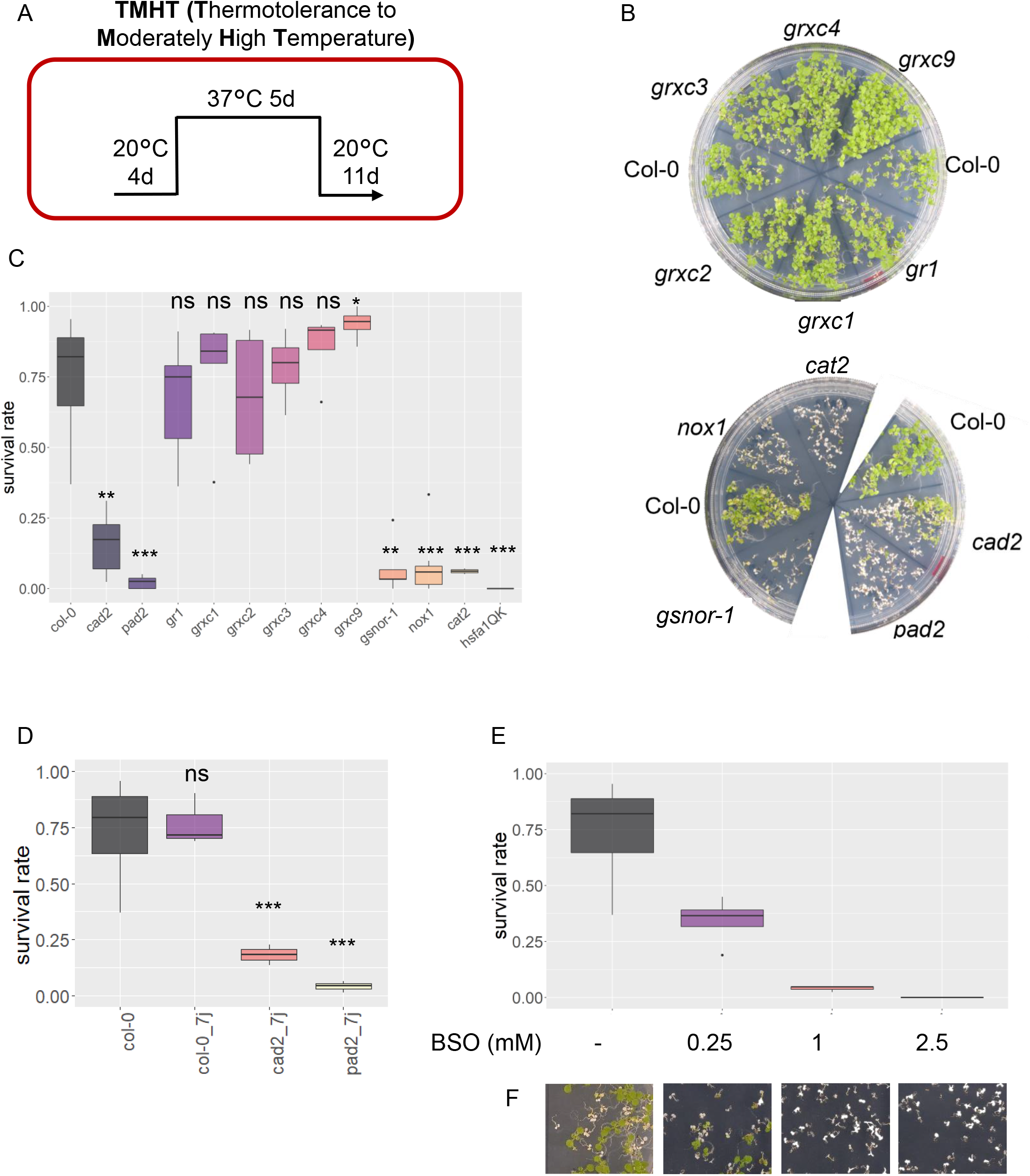
Low glutathione impairs thermotolerance. A-C, Wild-type plants (Col-0), *cad2, pad2, gr1, grxC1, grxC2, grxC3, grxC4, grxc9, gsnor-1, nox1, cat2* and Heat Shock Factor A1 quadruple knockout mutant (*hsfA1QK*) were subjected to a Thermotolerance to Moderate High Temperature (TMHT) regime. A, TMHT design. B, Plate pictures taken 11 days after recovery. C, Viability of the plants was assessed by the recovery of shoot growth after recovery. Data are means of at least 3 biological repetitions of 30 to 300 plants for each condition. Asterisks indicate significant differences compared to Col-0 by Student’s t test (P<0.01 = *, P<0.001 = **, and P<0.0001 = ***). D, Pictures of 7 days-old col-0, *cad2* and *pad2* plants after TMHT assay. E, Survival rate of col-0 plants grown on increasing GSH synthesis inhibitor concentration BSO (buthionine sulfoximine). F, pictures of the plates after recovery.

### Low glutathione impairs thermomorphogenesis

As we found that the glutathione steady-state levels are affected by a 27°C treatment, we also studied the behavior of glutathione deficient mutants to a 27°C high temperature treatment applied for 7 days (Figure 5A). As previously shown, this regime induces a thermomorphogenesis adaptation monitored by elongation of the hypocotyl (Figure 5). At 20°C, hypocotyl length of *cad2* and *pad2* mutants is like Col-0 (Figure 5B-C). However, when subjected to 27°C, the hypocotyl elongation observed in wild-type plants is partially impaired in both *cad2* and *pad2*, suggesting that low glutathione impairs high temperature-induced hypocotyl elongation. By observing the plants, we also noticed less elongated petioles in both mutants, which is another hallmark of thermomorphogenesis (Figure 5B). In contrast, high temperature-dependent hypocotyl elongation was not affected in the cytosolic glutathione reductase mutant (*gr1*), suggesting that glutathione level rather than glutathione redox state is involved in thermomorphogenesis (Figure 5C). As for TMHT, we also subjected grx mutants to the thermomorphogenesis regime. None of these mutants (excepted *grxC3* and *grxC4*) had a perturbed hypocotyl elongation phenotype, suggesting that most of these candidates are not involved in the glutathione deficient mutant’s phenotype (Figure 5D).

**Figure 5:**
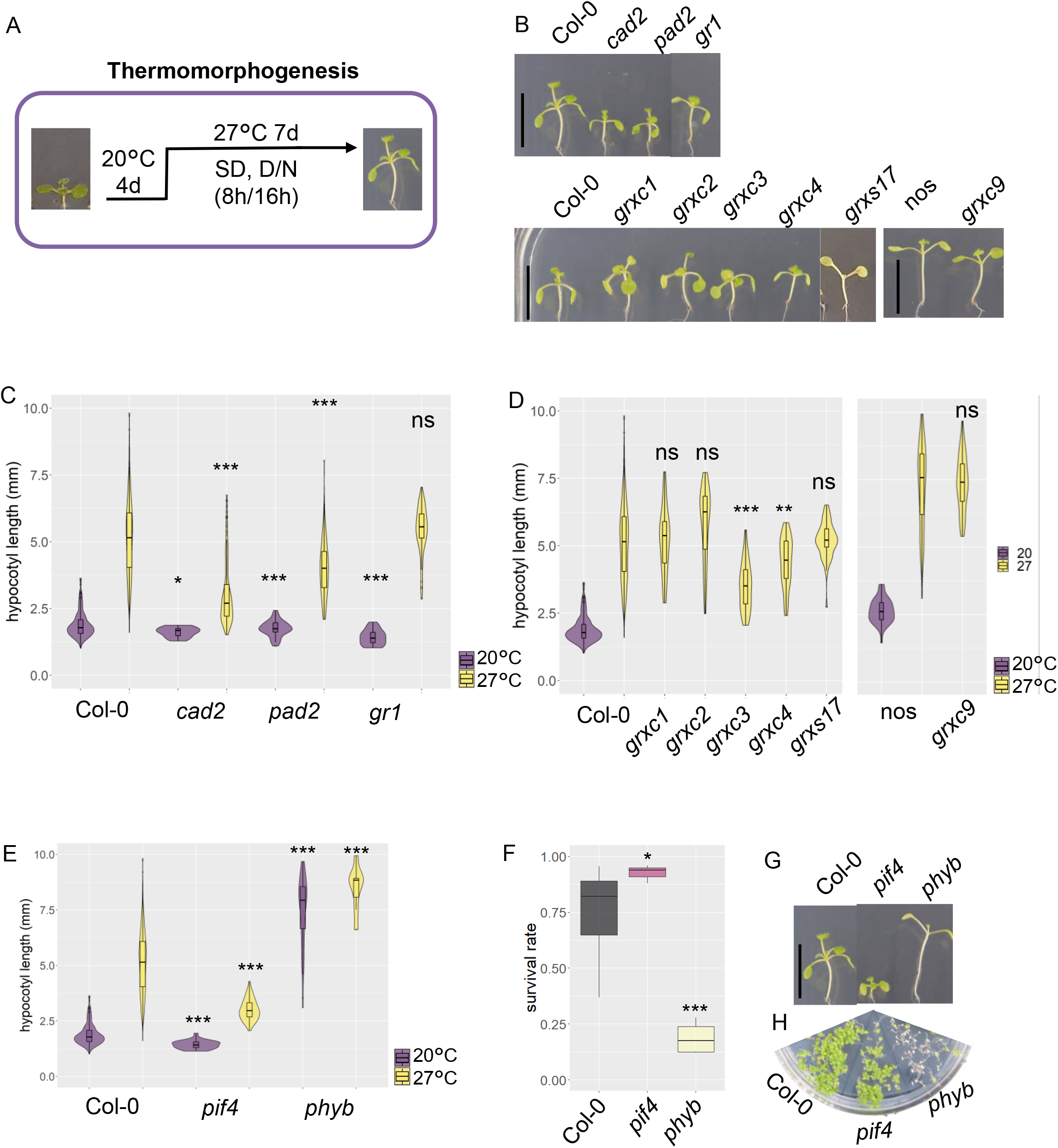
Low glutathione impairs thermomorphogenesis. A-C, Wild-type plants (Col-0 and nos), *cad2, pad2, gr1, grxC1, grxC2, grxC3, grxC4, grxS17* and *grxc9* were subjected to a 27°C thermomorphogenesis assay. A, Thermomorphogenesis assay design. B, Pictures of representative plants taken after 11 days treatment black bar = 10mm. C, D Hypocotyl length measurements. Pictures of plants were taken, and hypocotyl length was measured using the neuron plugging from ImageJ software. Data are means of at least 30 hypocotyls per condition. Asterisks indicate significant differences compared to Col-0 at 20°C or 27°C by Student’s t test (P<0.01 = *, P<0.001 = **, and P<0.0001 = ***). Phenotypes of *pif4* and *phyb* after thermomorphogenesis assay (E) and thermotolerance assay (TMHT) (F). Pictures of representative *pif4* and *phyb* plants after thermomorphogenesis (G) and TMHT (H).

Finally, we re-analyzed the hypocotyl elongation of two mutants (*pif4* and *phyB*) involved in the thermomorphogenesis regulation (Jung et al., 2016; Legris et al., 2016; Gangappa and Kumar, 2017). As previously described, both mutants have contrasted hypocotyl elongation phenotypes. While the *phyB* mutant has a long hypocotyl at both 20°C and 27°C, due to constitutively induced thermomorphogenesis, the *pif4* mutant fails to induce hypocotyl elongation at 27°C due to its impaired thermomorphogenesis pathway (Figure 5E). To study whether the lack of thermomorphogenesis of the *pif4* mutant also affects its thermotolerance, we subjected *pif4* to TMHT. The viability of the pif4 mutant is not affected, indicating that thermomorphogenesis pathway does not overlap with the tolerance to TMHT. However, the *phyB* mutant shows a survival rate to TMHT, which can be due to his long hypocotyl phenotype (Figure 5F-G).

To examine if glutathione is involved in the PHYB/PIF4-dependent thermomorphogenesis regulation, we applied low concentrations (0.2 mM) of exogenous BSO *on pif4* and *phyB* mutants and measured hypocotyl length under 20°C or 27°C. *pif4* plants treated with 0.2 mM BSO show reduced hypocotyl elongation at 27°C, suggesting another PIF4-independent thermomorphogenesis pathway that requires normal GSH level (Supplemental Figure 3). In the contrary, the *phyb* long hypocotyl phenotype is exacerbate by BSO treatment (Supplemental Figure 4).

### Low glutathione impairs transcriptional response to high temperature

As we observed a transient glutathione decrease in both cytosol and nuclear compartments in response to heat stress and that the response to heat is impaired in glutathione deficient mutants, we wished to explore the impact of both heat stress regimes on genome expression in both wildtype and the glutathione deficient mutant *cad2*. Therefore, we performed a wide transcriptomic analysis by RNA-seq. Wild-type and *cad2* plants were treated in the same conditions as previously described (20°C, 27°C and 37°C) and samples from three biological repetitions were taken before and during temperature treatments, at times coinciding with the decrease (2 h) and during the recovery of glutathione level (24 h) (Figure 2). The reliability of the RNAseq analyses was assessed by PCA analyses (Supplemental Figure 5) and differentially expressed genes were selected with a 2-fold cutoff (Padj<0.01) (Supplemental Dataset 1). We first analyzed the impact of heat stress on gene expression deviation in wild-type plants (Supplemental Figure 6). As expected, a 37°C treatment has a much stronger effect on gene expression reprogramming as a 27°C treatment: 12189 (5600 upregulated, 6499 downregulated) and 891 (437 upregulated, 454 downregulated) genes whose expression is modified after stress, respectively. A large majority of differentially expressed genes (64% up-, 90% down-regulated) at 27°C treatment was common with 37°C treatments, indicating that both treatments trigger a common gene regulation signature (Supplemental Figure 6C-D). As expected, gene ontology (GO) analyses associate common responses with ‘stimulus, stress and high temperature’ responses (Supplemental Figure 6B).

Then we evaluated the impact of the *cad2* mutation on gene expression after heat stress (Figures 6 and 7). At 27°C, we identified 387 genes that were differentially expressed upon high temperature in both genotypes. Consistent with the temperature treatment imposed to the plants, the shared regulatory signatures were associated with heat response [GO categories: response to heat, temperature, stress] (Figure 6B). We also observed contrasting regulatory responses between wildtype and *cad2* mutants. Heatmap analyses delinearated misregulated genes in different clusters. Among them, large clusters correspond to genes that failed to be induced upon high temperature [GO categories: response to abiotic stress and temperature stimulus, response to ROS] (Figure 6B-D). Interestingly, among differentially expressed genes between wild-type and *cad2*, GO categories underline a differential response to oxidative stress (Figure 6B). Taken together, these results show that in addition to common core regulatory signatures, the *cad2* mutant display specific responses to elevated ambient temperature.

**Figure 6:**
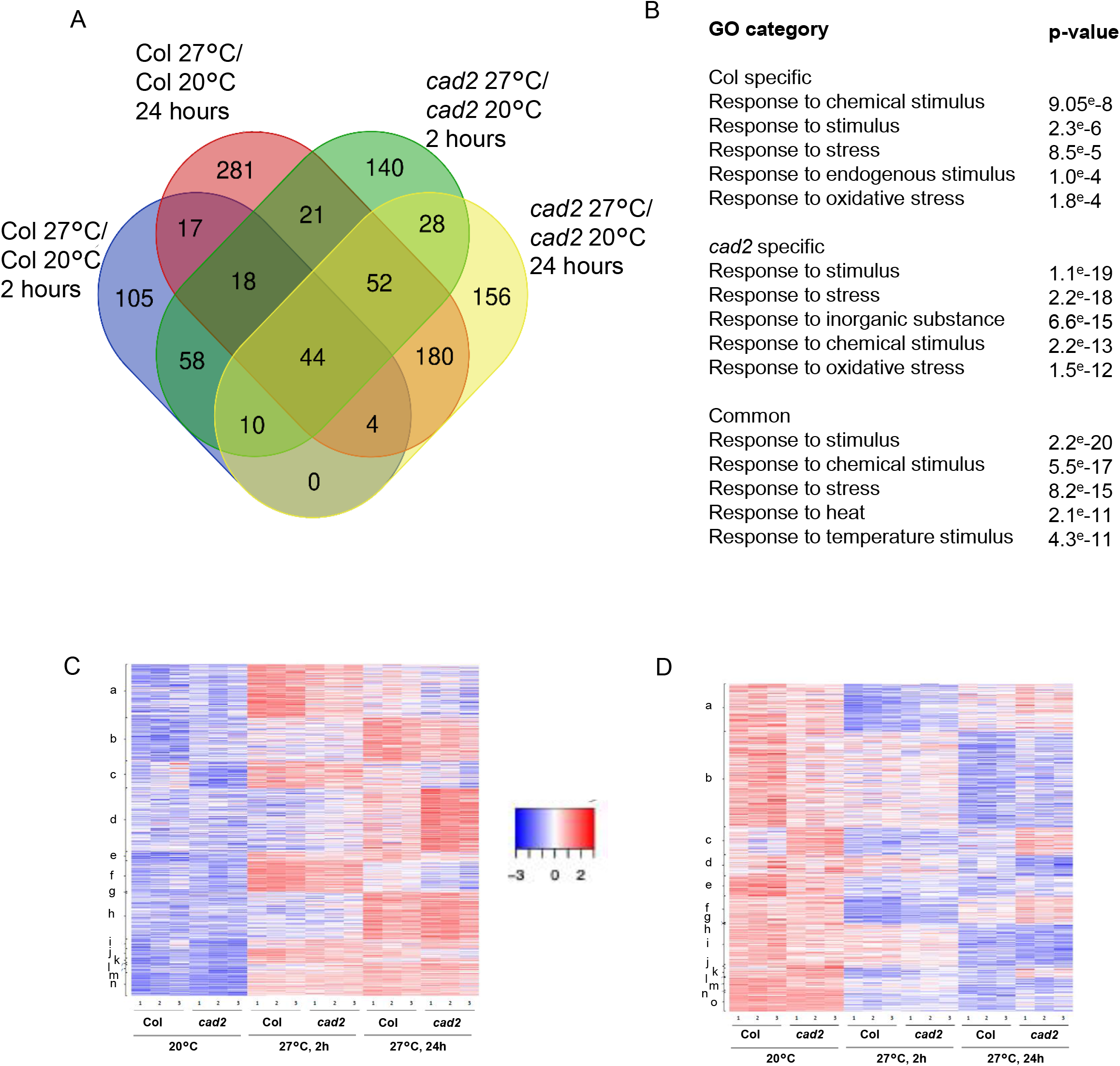
Genome-wide analysis of *cad2* response to 27°C temperature. A-B, Genes upregulated (A) or downregulated (B) at 2 h and 24 h (fold change cutoff log>1, Paj<0.05) after the temperature shift in wild-type (Col-0) and *cad2* mutant. Cutoff Gene ontologies (GO) characterize the biological processes enriched among the temperature-regulated genes that are shared or that are specific between the wild-type and the *cad2* mutant. C-D, Heatmaps of upregulated (A) or downregulated (B) genes in the three RNAseq biological repetitions (1,2,3) before and after high temperature shift. Genes were clustered as follows: a, Col-0 27°C 2h; b, Col-0 27°C 24h; c, *cad2 27°C* 2h; d, *cad2 27°C* 24h; e, common Col-0 27°C 2h and Col-0 27°C 24h; f, common Col-0 27°C 2h and *cad2 27°C* 2h; g, common Col-0 27°C 24h and *cad2 27°C* 2h; h, common Col-0 27°C 24h and *cad2 27°C* 24h; i, common *cad2 27°C* 2h and *cad2 27°C* 24h; j, common Col-0 27°C 2h, Col-0 27°C 24h and *cad2 27°C* 2h; k, Col-0 27°C 2h, *cad2 27°C* 2h and *cad2 27°C* 24h; l, common Col-0 27°C 2h, Col-0 27°C 24h and *cad2 27°C* 24h; m, common Col-0 27°C 24h, *cad2 27°C* 2h and *cad2 27°C* 24h; n, common for all.

**Figure 7:**
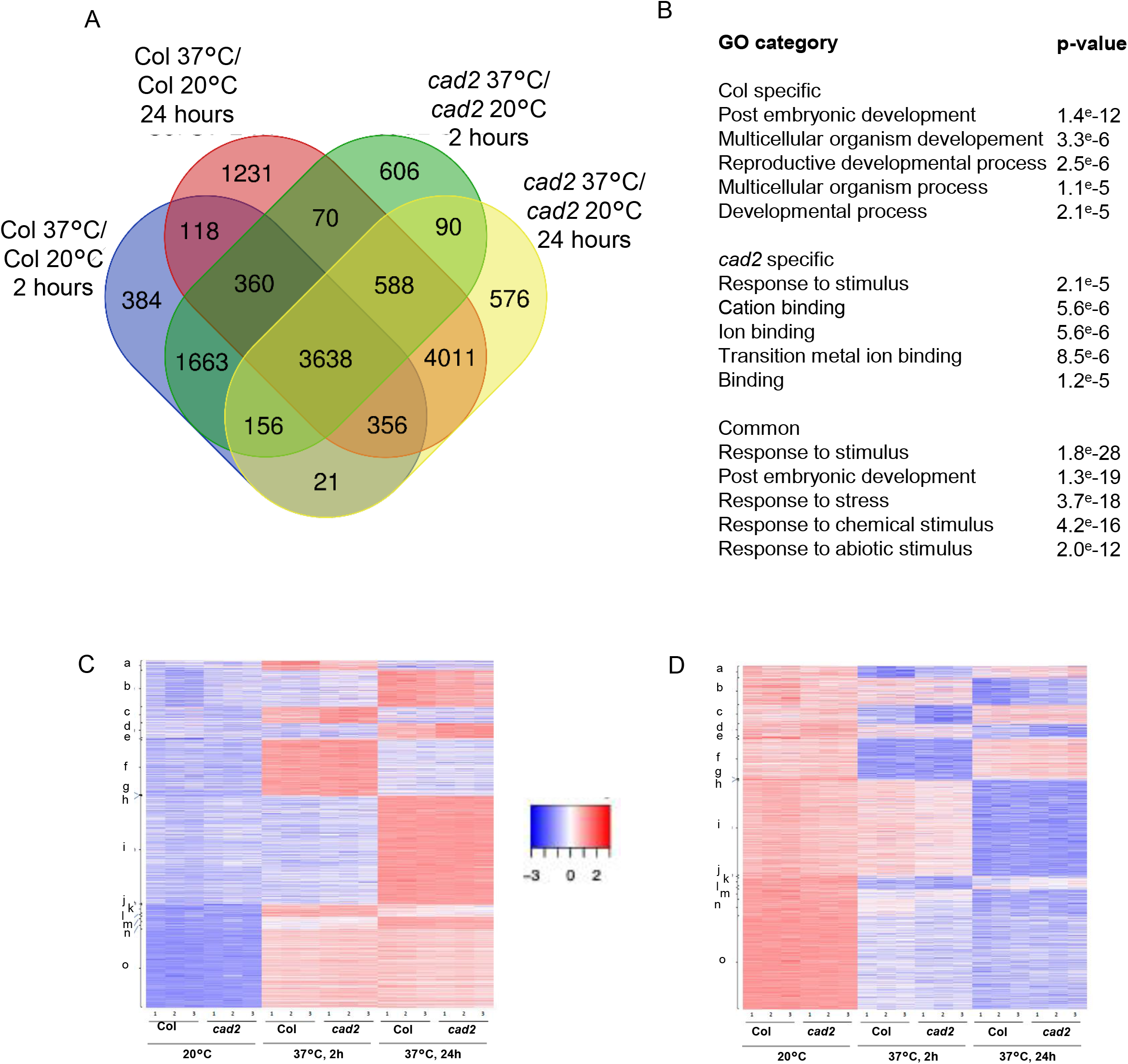
Genome-wide analysis of *cad2* response to 37°C temperature. A-B, Genes upregulated (A) or downregulated (B) at 2 h and 24 h (fold change cutoff log>1, Paj<0.05) after the temperature shift in wild-type (Col-0) and *cad2* mutant. Cutoff Gene ontologies (GO) characterize the biological processes enriched among the temperature-regulated genes that are shared or that are specific between the wild-type and the *cad2* mutant. C-D, Heatmaps of upregulated (A) or downregulated (B) genes in the three RNAseq biological repetitions (1,2,3) before and after high temperature shift. Genes were clustered as follows: a, Col-0 27°C 2h; b, Col-0 27°C 24h; c, *cad2 27°C* 2h; d, *cad2 27°C* 24h; e, common Col-0 27°C 2h and Col-0 27°C 24h; f, common Col-0 27°C 2h and *cad2 27°C* 2h; g, common Col-0 27°C 24h and *cad2* 27°C 2h; h, common Col-0 27°C 24h and *cad2* 27°C 24h; i, common *cad2* 27°C 2h and *cad2* 27°C 24h; j, common Col-0 27°C 2h, Col-0 27°C 24h and *cad2* 27°C 2h; k, Col-0 27°C 2h, *cad2* 27°C 2h and *cad2* 27°C 24h; l, common Col-0 27°C 2h, Col-0 27°C 24h and *cad2* 27°C 24h; m, common Col-0 27°C 24h, *cad2* 27°C 2h and *cad2* 27°C 24h; n, common for all.

Comparison of the gene expression profiles between wild-type and *cad2* plants subjected to 37°C also shows a large panel of common differentially expressed genes in both genetic backgrounds (78% of differentially expressed genes), indicating that major heat stress response pathways are not perturbed in the *cad2* mutant (Figure 7A-B). Among them, major heat stress response genes like HSPs (101, 90, 70, small HSPs) are normally expressed in the mutant, while differential expression were found in some HSF transcription factors (Supplemental Figure 7). However, major differentially expressed clusters were also found to be specifically expressed in wild-type or *cad2* mutants (Figure 7A). GO categories underline a differential response to developmental process in wild-type and to ion binding capacities in *cad2*, respectively (Figure 7B).

Finally, to characterize whether glutathione metabolism genes are differentially regulated by in the *cad2* mutant upon high temperature, we selected a panel of 140 genes encoding proteins involved in glutathione metabolism/catabolism or which use glutathione as substrate (Figure 8A and Supplemental Table 1) and analyzed their expression upon both high temperature regimes. Heatmap show that most of the genes show common response to high temperature, indicated that these genes are not much impacted by the stress treatments at the transcriptional level. Core glutathione synthesis (GSH1, GSH2) and reduction (GR1, GR2) do not fluctuate much in any mutant upon heat stress (Figure 8B). However, several genes encoding for glutathione-S-transferases (GSTs), show differential expression (Figure 8B). Among them, GSTU are known to be involved in S-glutathionylation which might be relevant for modulating glutathionylation upon heat stress.

**Figure 8:**
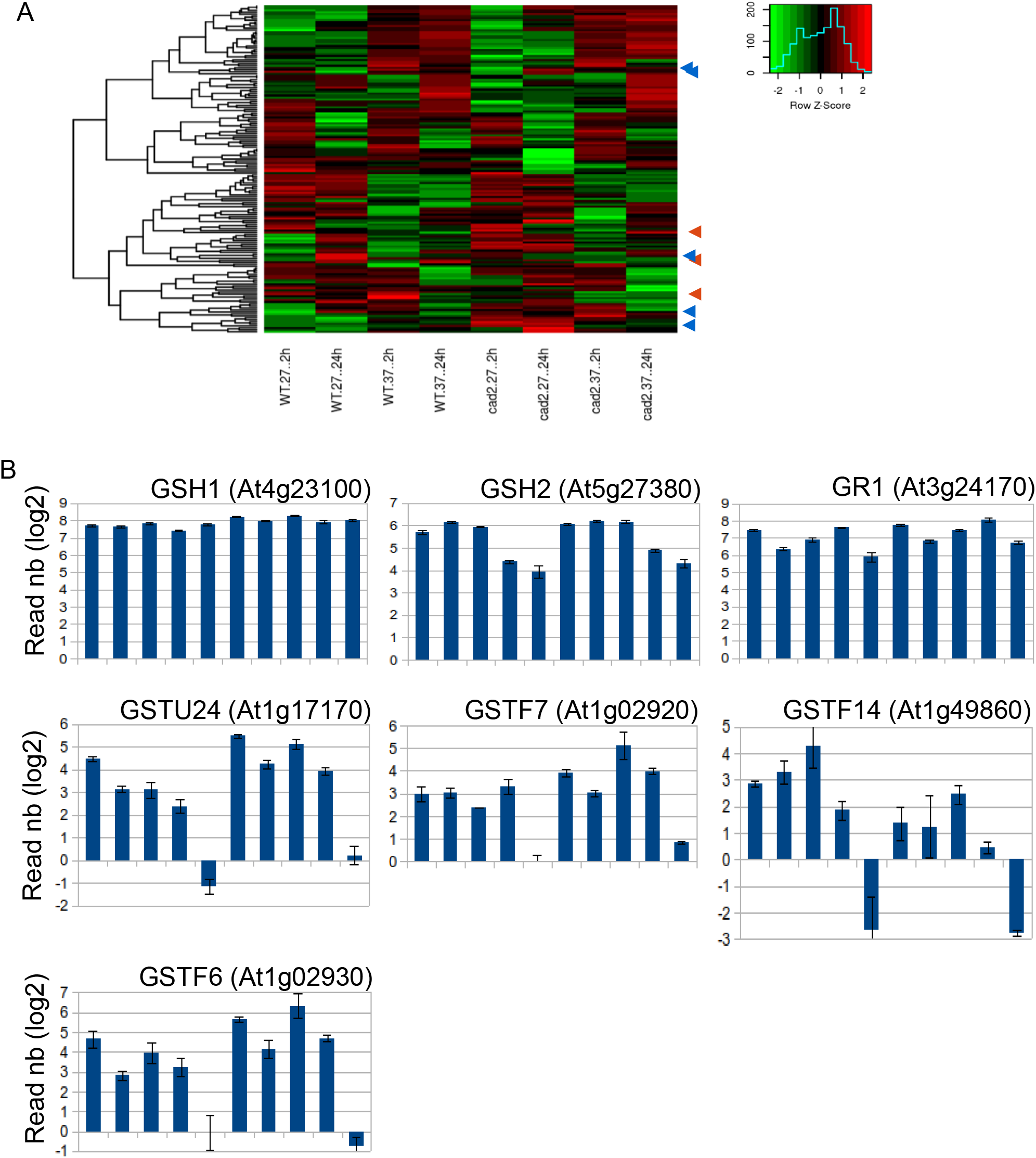
Glutathione-related gene expression upon high temperature. A, Heatmap of expression of glutathione related genes (shown in Supplemental Figure 1) under high temperature regimes, as compared to 20°C. Arrows highlight differentially expressed clusters at 27°C (blue) or at 37°C (orange) B, differentially expressed genes selected from RNAseq data.

All in all, the genome-wide gene expression analyses indicate, that while the glutathione deficient *cad2* mutant shares most common response to heat temperature with wild-type plants, it also shows deviations in the response which might participate to the sensitivity of the *cad2* mutant to both high temperature regimes.

## DISCUSSION

### High temperature leads to cytosolic and nuclear oxidation

During periods of heat stress, living organisms need to rapidly adapt to cope with the damaging effects of increasing temperatures. However, the mechanisms involved in the adaptation to heat stress largely differ depending on the duration and the intensity of the stress. In Arabidopsis, for which optimal temperature growth is generally established at 20-22°C, a moderate raise of the ambient temperature to 27°C leads to a change in plant morphogenesis called thermomorphogenesis, easily monitored by the elongation of the hypocotyl and involving different processes like gene expression reprogramming, chromatin remodeling or hormone accumulation (Casal and Balasubramanian, 2019). More intense raise of temperatures (e.g. 35-42°C) often found during summer heat waves are much more harmful for plant and damage cellular micromolecules like lipids and proteins. Accordingly, protecting mechanisms are rapidly induced, gene expression is massively reprogrammed and among them, protein chaperones are massively expressed. As heat stress is also known to induce accumulation of ROS and to modify the redox state of the cell, in this study, we investigated the redox modifications induced in two contrasted high-temperature regimes, 27°C inducing thermomorphogenesis and 37°C, triggering heat stress responses. Indeed, we show that, while a 27°C treatment hardly impact the redox state of the roGFP2, at least during the first hour after the treatment, a 37°C treatment rapidly induces a full roGFP2 oxidation (Figure 1). Indeed, a first estimation indicates that the glutathione redox potential decrease from −309mV +/-10 at 20°C to −247 mV +/-21 (Figure 1). In a previous report, Babbar et al. (2021) have also observed a marked change in the glutathione redox potential (about −330 mV to −250 mV) of the cytosol and nuclei of epidermal and stomatal guard cells after a 1h heat stress at 42°C. Consistently, our data support a similar oxidation of the roGFP in the nuclear compartment which seems to occur even faster as the cytosol at 37°C (Supplemental Figure 1). Such a slightly faster nuclear roGFP oxidation kinetic should be further confirmed by other way but could provide from a lower nuclear glutathione reduction capacity or a slight difference in the glutathione cytosolic/nuclear fluxes during heat stress (Diaz Vivancos et al., 2010; Delorme-Hinoux et al., 2016).

As a sensor of the redox state of glutathione, roGFP oxidation is generally associated with accumulation of oxidized glutathione, for example in glutathione reductases mutants or in plants accumulating high H_2_O_2_ levels (Marty et al., 2009; Marty et al., 2019). However, we did not observe a massive accumulation of oxidized glutathione after the raise of temperature but rather a strong decrease of the total glutathione pool (Figure 2). Although this decrease leads to a mild change of the GSSG/GSH ratio during the first hours after the treatments, such a slight modification of the GSSG/GSH ratio hardly explains the almost full oxidation of the roGFP. However, previous report shows that the roGFP redox state is not only dependent on the redox state of glutathione but also on variations in the glutathione pool. Pharmacological (BSO or CDNB) or genetic depletion of the free glutathione pool led to partial or total oxidation of the roGFP, depending on the extent of the depletion, even in conditions where the glutathione redox state is not affected (Meyer et al., 2007). Accordingly, roGFP has been found partially oxidized already at 20°C, in the *cad2* mutant which harbors a decreased glutathione level but no affected redox state of glutathione (Figure 3). Moreover, partial depletion of the glutathione likely further cause oxidation of the cell upon high temperature, due to a weakness of the antioxidant capacities, which we at least can observe by the accumulation of DAB staining in the root apex after heat stress (Supplemental Figure 2).

The reason for the decrease of the glutathione pool we observed after high temperature will have to be further determined. This might be partly due to conjugation to other molecules during heat stress through S-glutathionylation. Glutaredoxins, enzymes which catalyze S-glutathionylation, could be involved in the process, and many of them were identified as differentially expressed in our RNAseq analyses (Supplemental Table 1 and Dataset 1). Three other types of enzymes are involved in glutathione catabolism: carboxypeptidases, phytochelatin synthases, and gammaglutamyl transpeptidases (GGT) (Meister, 1988; Blum et al., 2007; Blum et al., 2010; Meyer et al., 2012). For example, analyses of Arabidopsis mutants have described functions of the GGT1 and GGT2 in preventing oxidative stress by metabolizing extracellular GSSG and transporting glutathione into developing seeds (Ohkama-Ohtsu et al., 2007). Our biochemical glutathione measurements also point to the capacity of plants to recover a steady state level of glutathione after three days of continuous high temperature treatment which likely participate to the adaptation of the plant to high temperature (Figure 2). While we do not observe increase of the glutathione biosynthesis gene (GSH1, GSH2) expression during the recovery period (Figure 8), we may assume that increased GSH1 or GSH2 enzyme activity might be responsible for this readjustment.

Therefore, we conclude that both moderate raise of ambient temperature (27°C) and a more intense high temperature increase (37°C) lead to perturbation of the glutathione pools rather than modification of the glutathione redox state, and that plant readjust a steady state level after a few days of these treatments.

### Adjusted glutathione pool is required for tolerance to high temperature regimes

The requirement of a “high” glutathione steady-state level to allow plant tolerance to high temperature is evidenced by the sensitivity of the glutathione deficient mutants (*cad2, pad2*) and BSO-treated plants, to high temperature regimes. Somehow supporting the minor change in the glutathione redox state upon high temperature, the *gr1* mutant inactivated in the cytosolic/nuclear glutathione reductase is hardly affected by high temperature (Figure 4 and 5). Nevertheless, the role of the thioredoxin pathways as a backup for glutathione reduction has also to be considered in the context of the plant response to high temperature (Marty et al., 2009; Marty et al., 2019).

The fact that the two high temperature regimes we studied here induce different plant responses imply that glutathione can be involved in both pathways (Figures 4, 5 and Supplemental Figure 5). Indeed, thermomorphogenesis induction involves many regulatory pathways which might be under redox control. The PHYB/PIF4 module play major role in sensing and transmitting the temperature signal to regulate gene expression. Among the induced pathways, hormones like auxin, brassinosteroids and GA are clearly involved. For example, hypocotyl growth and leaf hyponastic responses to warm temperatures are impaired in mutants deficient in auxin synthesis, perception or signaling (Gray et al., 1998; Sun et al., 2012). While our RNAseq analysis fail to identify clear GO categories of thermomorphogenesis genes among the differentially expressed genes in the *cad2* mutant, we do observe some differences in individual genes like *PIF4* or some auxin metabolism genes (Supplemental Figure 6). Moreover, connections between glutathione and auxin metabolism have been previously described in other plant development aspects (Bashandy et al., 2010; Schnaubelt et al., 2015; Trujillo-Hernandez et al., 2020). Also, glutathione-dependent glutaredoxins are suggested to regulate brassinosteroids pathway (Bender et al., 2015).

In a similar way, plant tolerance to the TMHT regime (37°C) requires an appropriate pool of glutathione, as both *cad2, pad2* and BSO-treated plants are unable to survive to the treatment (Figure 5). However, an affected glutathione redox state in the *gr1* seems not a major factor for the survival with respect of alternative reduction pathway occurring in the cytosol and the nucleus (Marty et al., 2009).

### High temperature transcriptional reprogramming is partially glutathione-dependent

In line with a role of glutathione in thermotolerance, our genome-wide gene expression analyses identified substantial deviations in gene expression reprogramming in the *cad2* mutant (Figures 6-8), suggesting that the transcriptional responses to high temperatures are partially altered in the mutant. To a certain extent, our data are consistent with previous genome-wide gene expression analyses in other glutathione deficient mutants (Ball et al., 2004; Han et al., 2013; Schnaubelt et al., 2015). A role of glutathione in gene expression regulation is also supported by the rapid roGFP oxidation we found in the nucleus under high temperature (Supplemental Figure 1). Although major high temperature response pathway (e.g. HSPs) were found to be induced as in the wild-type, large clusters of genes appeared to be much less induced in the mutant (Figure 6 and 7). Among the GO categories identified, we can assume that some “high temperature/stress response” pathways are misexpressed. Nevertheless, this does not mean that this transcriptional deviation is responsible to the thermotolerance defect of the *cad2* mutant. For example, the observed decrease in the expression of PIF4 in the mutant subjected to 27°C can be the consequence of impaired thermomorphogenesis, rather than its cause (Supplemental Figure 5). How glutathione is acting in the thermomorphogenesis pathways will have to be further studied. The observation that some glutaredoxins mutants also harbor hypocotyl elongation defects gives further clues for future research (Figure 5).

When focusing our RNAseq analyses on glutathione-dependent genes, it is interesting to notice that many GSTs have differential expression levels after high temperature. This is more pronounced under the 37°C regime, for which protein S-glutathionylation was found (Supplemental Figure 8), suggesting that proteins involved in glutathione conjugation might be particularly mobilized. In contrast, a batch of glutaredoxins, which generally act in protein S-deglutathionylation seems downregulated upon heat stress. However, no high temperature sensitivity was noticed in most of the glutaredoxin mutants we analyzed. This might be due to redundancy between members of the large glutaredoxins family (Riondet et al., 2012). However, an impaired thermotolerance of the *grxS17* was previously reported (Cheng et al., 2011; Knuesting et al., 2015; Martins et al., 2020a). While at 27°C, the *grxS17* mutation was suggested to impair auxin signaling (Cheng et al., 2011), the 35°C treatment was also suggested to induce holdase activities of GRXS17 (Martins et al., 2020a). Whether a lower glutathione level would limit GRXS17 thermotolerance activities should be further investigated.

## Conclusions

In this study, we describe the dynamics of redox changes occurring under two different high temperature regimes. Using GRX1:roGFP2 redox sensor, we show that high temperature induces oxidation of the cytosolic and nuclear compartment, which is mainly due to a decrease of the reduced glutathione level. However, plants can restore steady state level after at least one day of treatment, such adaptation likely being required both for the temperature-dependent developmental (thermomorphogenesis) and tolerance capacities of the plant. Further investigations are required to decipher the downstream actors of the glutathione-dependent adaptation to high temperature, but our genome-wide transcriptomic data suggest that genome reprogramming likely play a key role in this response.

## METHODS

### Plant materials and growth conditions

Wild type (WT) *Arabidopsis thaliana* ecotype Columbia-0 (Col-0) and Nossen_0 (Nos_0) ecotypes and different mutants were used for the experiments. The following mutants were all previously published and available in our team: *cad2-1* (*Howden et al., 1995*), *pad2-1* (Parisy et al., 2007), *gr1-1* (Marty et al., 2009), *cat2-1* (Queval et al., 2007), *gsnor/hot5-2* (Lee et al., 2008), *hsfA1QK* (Yeh et al., 2012), *grxC1* (Riondet et al., 2012), *grxC2* (Riondet et al., 2012), *grxC9* (Huang et al., 2016), *pif4-2* (Koini et al., 2009), phyb-9 (Reed et al., 1993). *grxC3* and *grxC4* (Salk_062448 and Salk_128264) were ordered from the Nottingham Arabidopsis Stock Centre (NASC; http://arabidopsis.info/).

For *in-vitro* plant culture, Arabidopsis seeds were surface-sterilized for 15 min with ethanol 70%, rinsed with ethanol 95% and completely dried before used. Sterilized seeds were plated onto ½ MS agar medium (2.2 g/L Murashige and Skoog, 0.5 g/L MES, 0.5% plant agar (w/v), adjusted to pH 5.8) added or not with buthionine sulfoximide (BSO, Sigma-Aldrich) and stratified 48 h at 4°C in the dark. Seedling were grown under 120 μmol.m^-2^. s^-1^ photosynthetic flux at 20°C and 16 h light/8 h dark cycles. Those growth conditions were systematically used, unless otherwise indicated.

### Thermotolerance assays

Seeds were first germinated at 20 °C, and after 4 to 7 days, seedlings were either transferred to 27 °C under short day (for hypocotyl length assay) or at 37°C under long day condition (for plant survival assay). Heat treatments were done by transferring plants in pre-heated MLR-352 Plant Growth Chambers (PHCbi, Sanyo). For thermomorphogenesis assays, pictures of each replicate were taken and hypocotyls of at least 20 seedlings were measured using NIH ImageJ software with the NeuronJ plugin. For TMHT experiment, the percentage of viability was calculated by counting the plants that recovered leaf growth 11 days after heat treatment. About 30 to 300 plants were used in each replicate for calculation and mean were plotted ± SD of 3 independent biological replicates. Asterisks indicate a significant difference (P<0.01 = *, P<0.001 = **, and P< 0.0001 = ***) using Student’s *t*-test for pairwise comparison to the WT (Col-0).

### Samples preparation and RNA-seq analysis

Seven days old seedlings were heated at 27°C or 37°C for 2h 30 min or 24h and control samples (20°C) were harvested just before the heat treatment. To take account of the contribution of time of day and the circadian clock to the heat stress responsive transcriptome in Arabidopsis treatments were done in the early morning, after 3h30 exposure to light (Blair et al., 2019). For each condition 3 biological replicates corresponding to different batch of plants were harvested and treated independently.

Each replicate was ground in liquid nitrogen with mortar and pestle and 100 mg of powder was used for RNA extraction using the Monarch^®^ Total RNA Miniprep Kit. Briefly, powder was suspended in RNA protection reagent, vortexed and centrifuged 2min (16,000 x g). An equal volume of RNA Lysis buffer was mixed with supernatant, loaded in the gDNA removal column in the 2min (16,000 x g) and the flow through wax mixed with equal volume of 100% ethanol. The mixture was then loaded into the RNA purification column, spin 30s and incubated with DNase 1 for 15 min and washed with RNA priming buffer, RNA Wash buffer two times and RNA was eluted in 70 μl of nuclease-free water and kept at −80 °C. Next, 2 μg of total RNA was purified with Dynabeads Oligo-dT (25) and library quality was determined using Bio-Analyser 2100 (Agilent Genomics). Quantification was done using Q-BIT Qubit Fluorescence Reader (ThermoFisher Scientific) and replicates were then multiplexed with unique barcodes. Library amplification, RNA-sequencing and data analysis for each mRNA library, single-end 75 base pair sequences were sequenced using NextSeq 550 instrument (Illumina) at the Bio-environment platform, University of Perpignan Via Domitia (UPVD). Reads were trimmed using Trimmomatic (Bolger et al., 2014), and mapped to the A. thaliana genome (Arabidopsis TAIR10 genome) using HISAT2 (Kim et al., 2015). The sequence alignment files were sorted and indexed using SAMtools (Li et al., 2009). Number of reads mapped onto a gene were calculated using HTSeq-count (Anders et al., 2015). Differentially expressed genes were obtained with DESeq2 (Love et al., 2014), using a log2 fold-change >1 (upregulated genes) or < −1 (down-regulated genes) with an adjusted *p*-value of 0.01. PCA were realized using PCA function from R. Heat map were visualised using the heatmap2 function from the R gplots package.

### Protein extraction and glutathionylation analysis

For total protein extraction, about 1 g of 7 days-old seedlings were ground in liquid nitrogen with mortar and pestle and suspended in 250 μL of protein extraction buffer (25 mM Tris pH 7.6, 75 mM NaCl, 0.1 % NP40 with one tab/10 ml of complete protease inhibitor cocktail (Roche). The solution was vortexed, centrifuged (13 000 g, 20 min, 4 °C) and the supernatant was taken up and the protein concentration was determined using the Bradford assay (Biorad).

Proteins were heated for 3 min at 95 °C in 0.6X Leammli buffer (62.5 mM Tris-HCl (tris(hydroxyméthyl) aminométhane) pH 6.8, 2.3 % SDS, 10 % glycerol, 0.01 % bromophenol blue). 25μg of protein extract was on Sodium Dodecyl Sulfate Polyacrylamide Gel Electrophoresis (SDS-PAGE) in 10% acrylamide gel (10 % acrylamide, 70 μM TEMED, 0.0004 % ammonium persulfate, 0.376 M Tris-HCl pH 6.3, 0.001 % SDS and miliQ water). The gel was transferred to Polyvinylidene difluoride (PVDF) membrane (90 min,0.16 mA), the membrane was then incubated for 1 h in a blocking solution composed of 0.5 % semi-skimmed milk powder in TBST (Tris-buffered saline 20 mM, 0.1 % Tween 20) and then incubated with the anti-glutathione (Invitrogen) primary antibody solution at 4 °C overnight. After washing with TBST (3×5 min), the membrane was incubated 3 h with a solution containing a secondary goat-HRP anti rabbit (1:10 000) (Biorad) (milk 0.5 % TBST). After washing with TBST (3×5 min), the revelation was carried out by incubating the membrane using the substrate HRP immobilon Western kit.

### Detection of reactive oxygen species

Detection of ROS was performed as previously described (Mhamdi et al., 2010). Staining was performed at room temperature on Col-0 7 days-old plants. For the detection of H_2_O_2_, plants were vacuum infiltrated in the Nitro Blue Tetrazolium (NBT) staining medium (10 mM phosphate buffer pH 7.5, 10 mM Na azide, one tablet of NBT (10 mg Sigma)). For the detection of superoxide, plants were vacuum infiltrated in 5 mM 3, 3’-Diaminobenzidine (DAB) at pH 3.8. For both staining, plants were then incubated in the same medium until the coloration was observed and then chlorophyll was removed in 95% ethanol before pictures taking.

### Glutathione measurements

Total glutathione content of 8-day-old seedlings was determined using the recycling enzymatic assay (Rahman et al., 2006). This method involves the reduction of GSSG by glutathione reductase and NADPH to GSH. GSH levels are determined by its oxidation by 5,5⍰-dithio-bis (2-nitrobenzoic acid) (DTNB), that produces a yellow compound 5⍰-thio-2-nitrobenzoic acid (TNB), measurable at 412 nm (Rahman et al., 2006). Briefly, 100 mg of fresh plant material ground in liquid nitrogen was extracted in 0.5 ml of 0.1 M Na phosphate buffer, pH 7.6, and 5 mM EDTA. After microcentrifugation (10 min, 9000 g), total glutathione in 0.1 ml of the supernatant was measured by spectrophotometry in a 1 ml mixture containing 6 mM DTNB (Sigma-Aldrich), 3 mM NADPH, and 2 U of glutathione reductase from Saccharomyces cerevisiae (Sigma-Aldrich). Glutathione-dependent reduction of DTNB was followed at 412 nm. The total glutathione level was calculated with the equation of the linear regression obtained from a standard GSH curve. GSSG was determined in the same extracts after derivatization of reduced GSH. Derivatization of 100 ml of plant extract was performed in 0.5 ml of 0.5 M K phosphate buffer, pH 7.6, in the presence of 4 ml of 2-vinylpyridine (Sigma-Aldrich) during 1 h at room temperature. The GSH-conjugated 2-vinylpyridine was extracted with 1 vol. of diethylether, GSSG was measured by spectrophotometry as described for total glutathione.

### Confocal Laser-Scanning Microscopy and Ro-GFP2 imaging

Observations were done using the Axio observer Z1 microscope with the LSM 700 scanning module and the ZEN 2010 software (Zeiss). Excitation of roGFP2 was performed at 488 and 405 nm and a bandpass (BP 490-555 nm) emission filter was used to collect roGFP2 signal. For background subtraction, signal was recovered using a BP 420-480 nm emission filter during excitation at 405 nm. Analysis and quantifications were performed as previously described (Schwarzländer et al., 2008), using the program ImageJ 1.52i (https://imagej.nih.gov/ij/).

## AUTHOR CONTRIBUTION

AD, CR and JPR designed research; AD, LB and AW performed research; NP and FP contributed new reagents/analytic tools; AD, NP, FP CR and JPR analyzed data; and CR and JPR drafted the paper.

## ACKNOWLEDGMENTS

This work was supported by the Centre National de la Recherche Scientifique, by the Agence Nationale de la Recherche (grant nos. ANR-REPHARE 19–CE12–0027 and ANR-RoxRNase 20-CE12-0025). This project was funded by Labex AGRO (under I-Site Muse framework) and coordinated by the Agropolis Fondation (grant no. Flagship Project 1802-002 - CalClim). This study is set within the framework of the Laboratoires d’Excellence TULIP (ANR-10-LABX-41). AD is supported by a Ph.D. grant from the Université de Perpignan Via Domitia (Ecole Doctorale Energie et Environnement ED305). We are grateful to Michèle Laudié and Margot Doberva from the Bio-Environment platform (University of Perpignan Via Domitia) for technical support in library preparation and sequencing.

## SUPPLEMENTAL FIGURES LEGENDS

**Supplemental Figure 1: *In vivo* monitoring of the nuclear glutathione redox state upon heat stress.** Fluorescence ratio calculated from samples shown in figure 1. ROI containing nuclei (gray bars), or cytosol (black bars) were selected from confocal images of cotyledon cells of ten-day old Arabidopsis wild-type seedlings stably expressing the Grx1-roGFP2 construct and subjected to 37°C. roGFP2 fluorescence was collected at 505-530 nm after excitation with either 405 nm or 488 nm. Ratio images were calculated as the 405/488 nm fluorescence. To fully reduced or oxidize the sensor, seedlings were immersed in 10 mM DTT or 100 mM H_2_O_2_, respectively. Control samples were immersed in MS/2 liquid medium and observed at 20°C. Then, the temperature of the thermostatic chamber was increased and the roGFP2 fluorescence were monitored 10 min for 1 hour. n=6.

**Supplemental Figure 2: ROS detection upon heat stress.** Four days old Col-0 seedlings were subjected to 37°C high temperature treatment during 2 hours before DAB and NBT staining. Representative pictures are shown. N=15 to 20.

**Supplemental Figure 3: Effect of BSO on *pif4* elongation at 27°C.** Four-days-old col-0 and *pif4* plants grown on 0.2 mM BSO were subjected to a thermomorphogenesis assays (as described in figure 4).

**Supplemental Figure 4: Effect of BSO on *phyb* elongation at 27°C.** Four-days-old col-0 and *phyb* plants grown on 0.2 mM BSO were subjected to a thermomorphogenesis assays (as described in figure 4).

**Supplemental Figure 5: PCA analysis of all RNAseq samples.**

Principal component analysis (PCA) performed for first two components of the thirty samples corresponding to the RNA-seq of three replicates for Col-0 and *cad2* mutant, at 27°C or 37°C for 2h or 24h and control samples (20°C).

**Supplemental Figure 6: Genome-wide analysis of wild-type response to 27°C and 37°C.** A-B, Genes upregulated (A) or downregulated (B) at 2 h and 24 h (fold change cutoff log>1, Paj<0.05) after the temperature shift in wild-type (Col-0). Cutoff Gene ontologies (GO) characterize the biological processes enriched among the temperature-regulated genes that are shared or that are specific between 27°C and 37°C. C-D, Heatmaps of upregulated (A) or downregulated (B) genes in the three RNAseq biological repetitions (1,2,3) before and after high temperature shift. Genes were clustered as follows: a, Col-0 27°C 2h; b, Col-0 27°C 24h; c, Col-0 37°C 2h; d, Col-0 27°C 24h; e, common Col-0 27°C 2h and Col-0 27°C 24h; f, common Col-0 27°C 2h and Col-0 37°C 2h; g, common Col-0 27°C 2h and Col-0 37°C 24h; h, common Col-0 27°C 24h and Col-0 37°C 2h; i, common Col-0 27°C 24h and Col-0 37°C 24h; j, common Col-0 37°C 2h and Col-0 37°C 24h; k, common Col-0 27°C 2h, Col-0 27°C 24h and Col-0 37°C 2h; l, common Col-0 27°C 2h, Col-0 37°C 2h and Col-0 37°C 24h; m, common Col-0 27°C 2h, Col-0 27°C 24h and Col-0 37°C 24h; n, common Col-0 27°C 24h, Col-0 37°C 2h and Col-0 37°C 24h; o, common for all.

**Supplemental Figure 7: Expression data of selected genes under high temperature.** Differentially expressed genes selected from RNAseq data.

**Supplemental Figure 8: Global protein S-glutathionylation increases at 37°C.** A, Western-blot analysis of the S-glutathionylated proteins of total protein extract from Col-0 plants at 37°C. Total protein was extracted, run on SDS-PAGE, and used for Western-blot with antibody against S-SG (1:1000), Total protein staining with Coomassie blue is shown as loading control. B, Intensity of each line was quantify using ImageJ and mean +-SD of 3 replicates was shown (Student’s t test (P<0.05= *, P<0.01 = **, and P<0.001 = ***).

**Supplemental Dataset 1: Raw data of RNAseq analyses.**

**Supplemental Table 1: Selected glutathione-dependent genes and their expression data upon high temperature.**

