## Supplemental Figures for "Glutathione-mediated plant response to high-temperature"

Figure S1

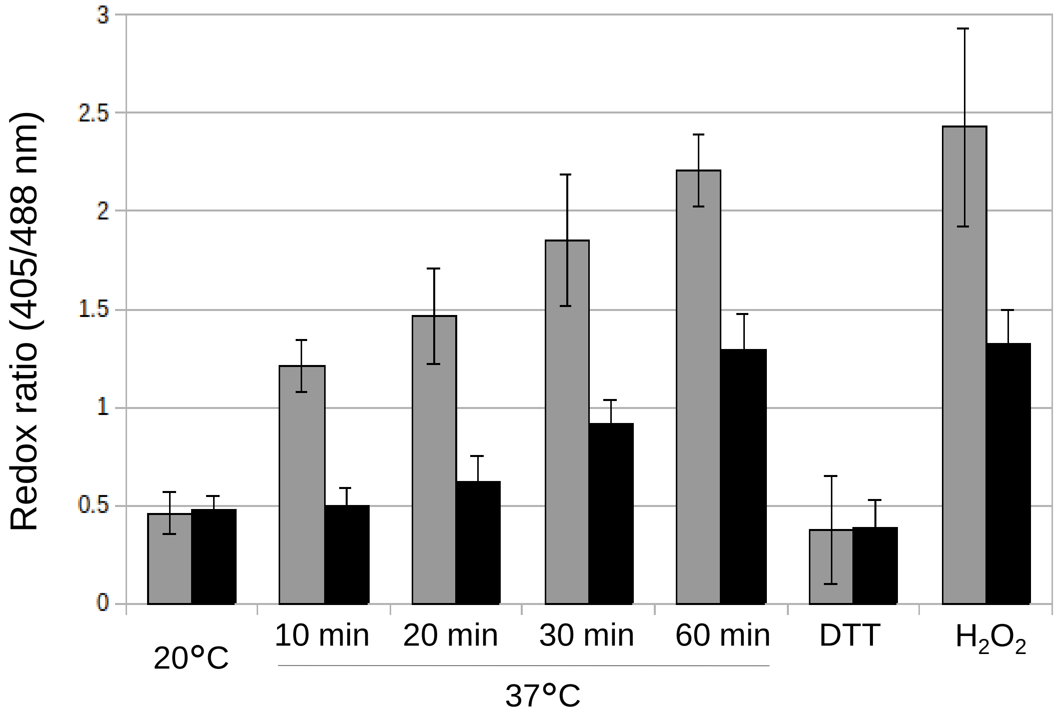
Redox ratio (405/488 nm)
3
2.5
2
1.5
1
0.5
0
20°C
10 min
20 min
30 min
60 min
DTT
$H_2O_2$
37°C

Figure S2

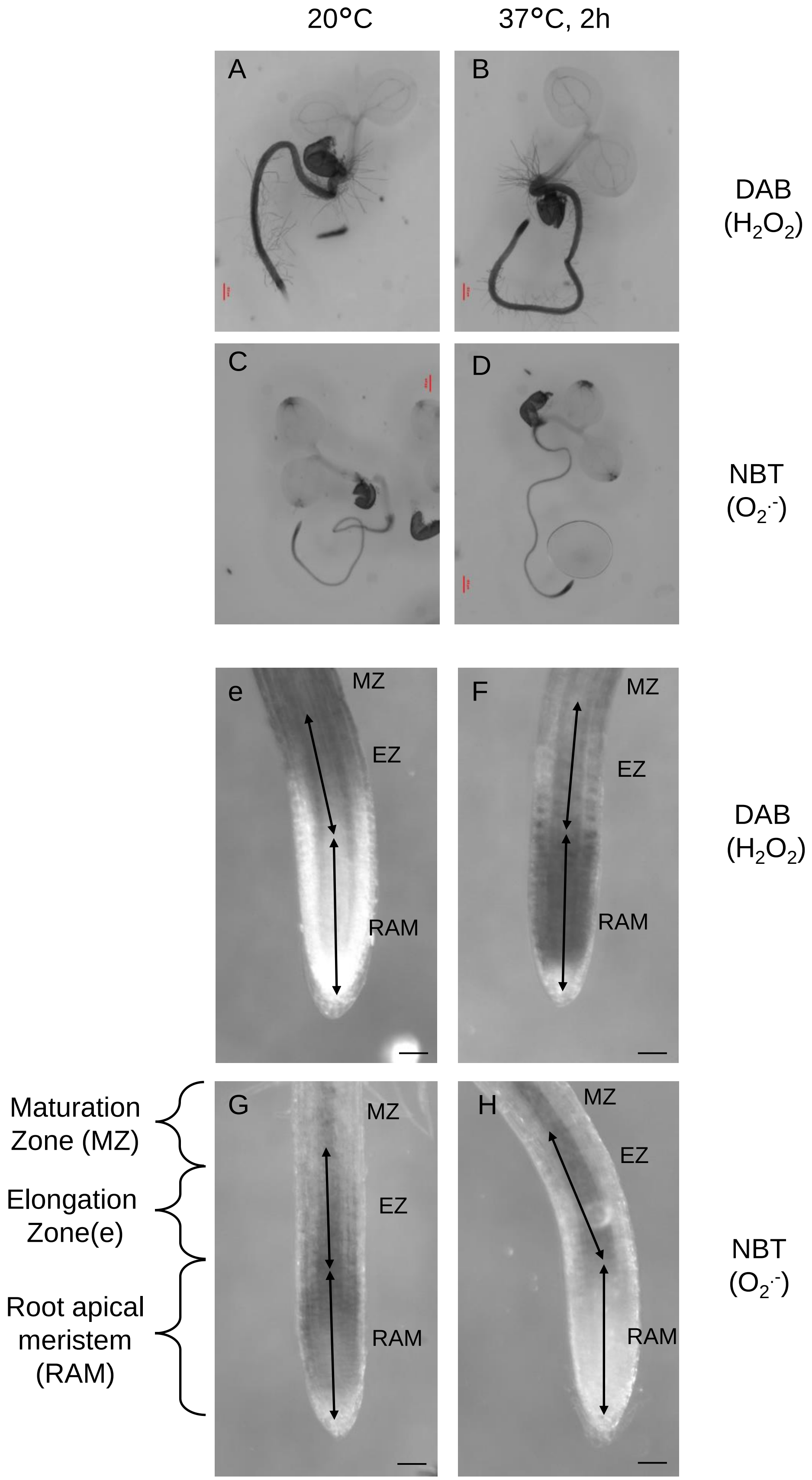

20°C
37°C, 2h
A
B
DAB
($H_2O_2$)
C
D
NBT
($O_2^{\cdot-}$)
e
MZ
EZ
RAM
F
MZ
EZ
RAM
DAB
($H_2O_2$)
Maturation
Zone (MZ)
Elongation
Zone(e)
Root apical
meristem
(RAM)
G
MZ
EZ
RAM
H
MZ
EZ
RAM
NBT
($O_2^{\cdot-}$)

Figure S3

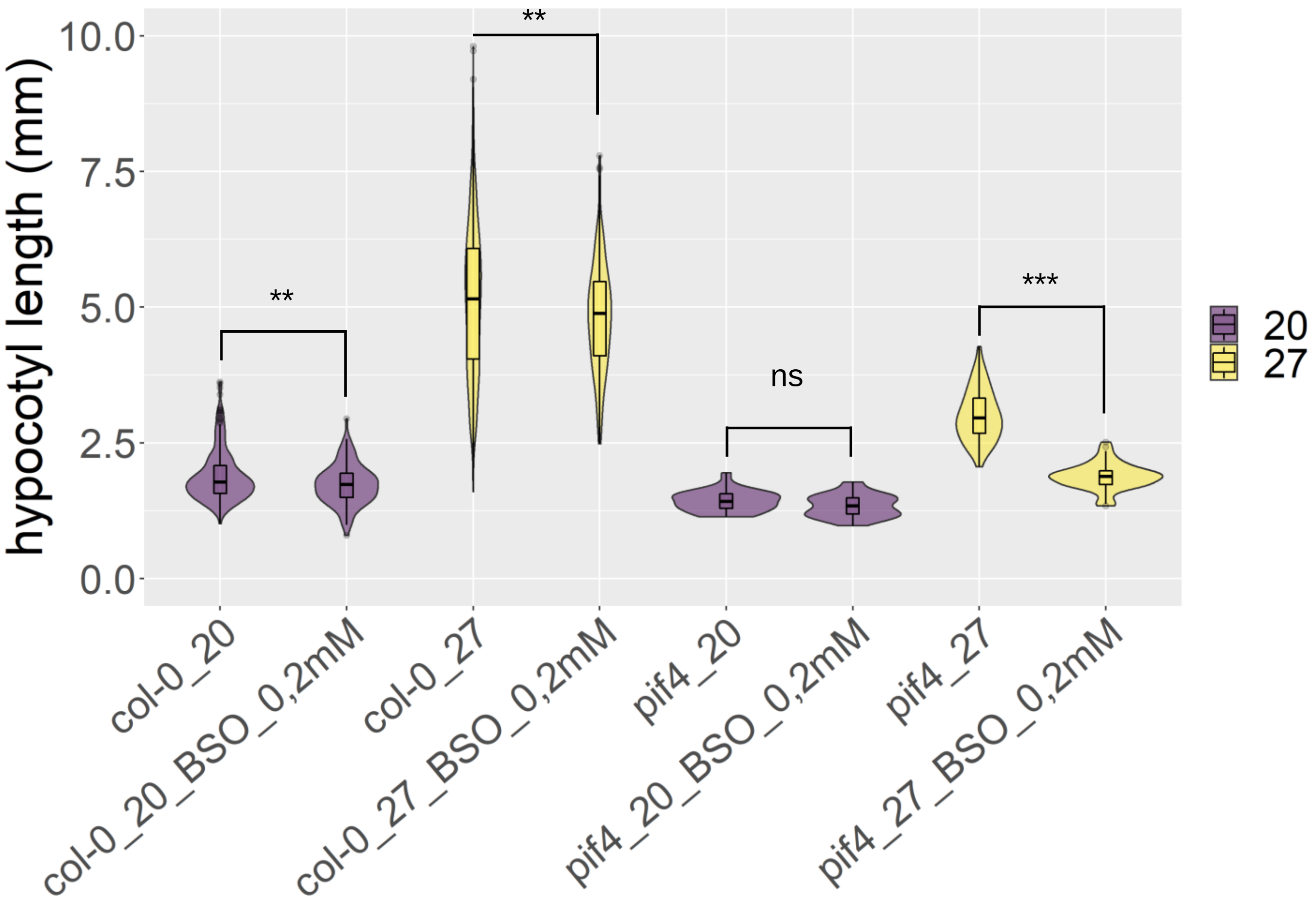

hypocotyl length (mm)
10.0
7.5
5.0
2.5
0.0
**
**
ns
***
20
27
col-0_20
col-0_20_BSO_0,2mM
col-0_27
col-0_27_BSO_0,2mM
pif4_20
pif4_20_BSO_0,2mM
pif4_27
pif4_27_BSO_0,2mM

Figure S4

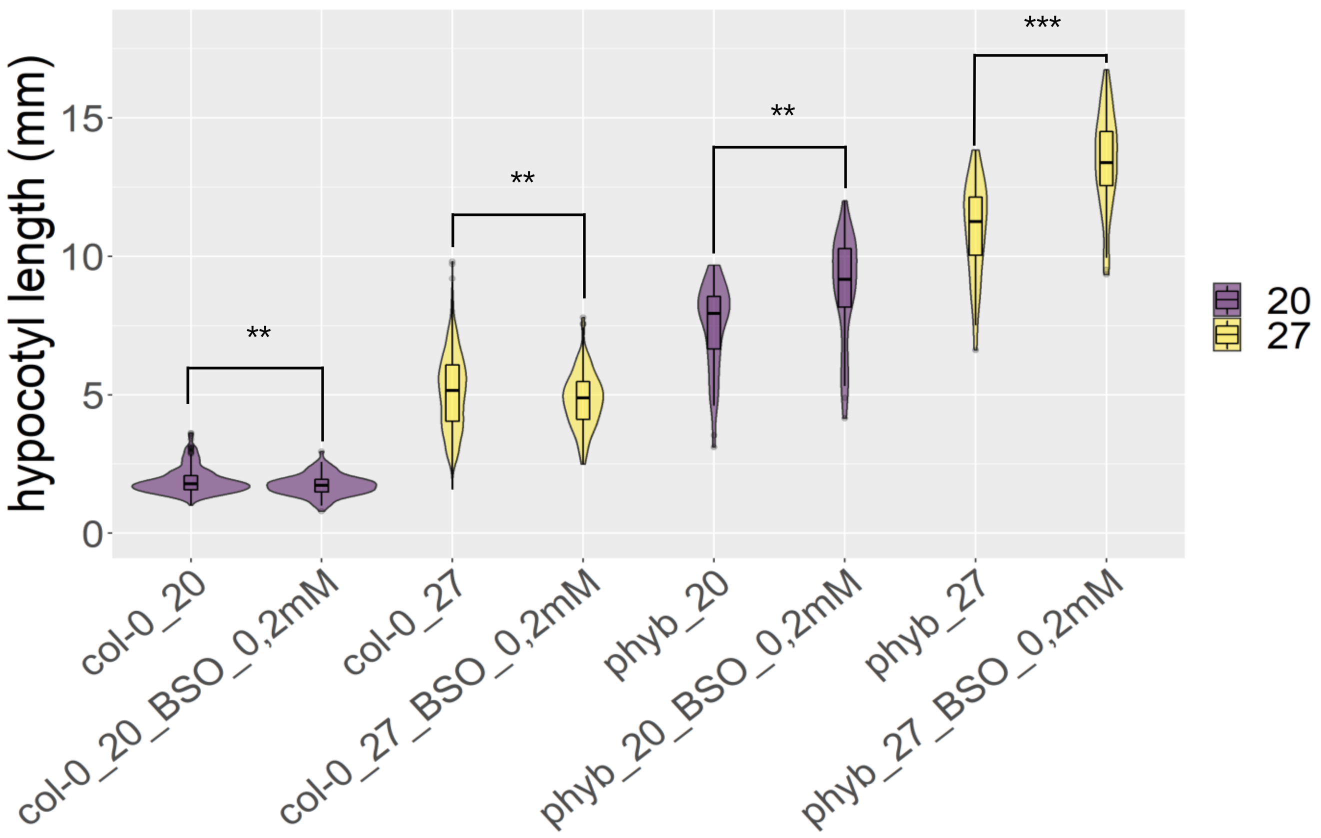

hypocotyl length (mm)
0
5
10
15
**
**
**
***
col-0_20
col-0_20_BSO_0,2mM
col-0_27
col-0_27_BSO_0,2mM
phyb_20
phyb_20_BSO_0,2mM
phyb_27
phyb_27_BSO_0,2mM
20
27

Figure S5

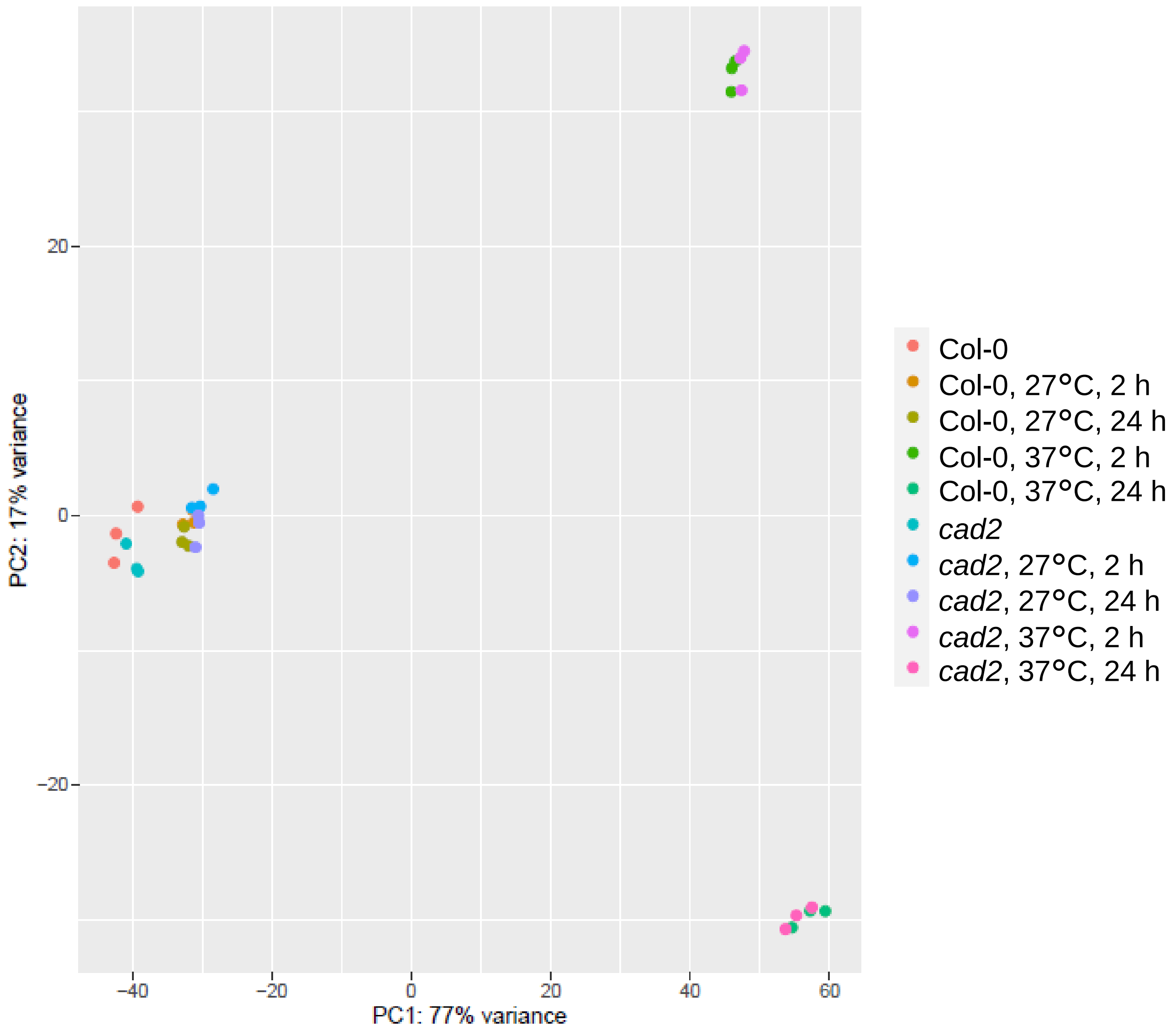

PC2: 17% variance
20
0
-20
-40
-20
0
20
40
60
PC1: 77% variance
Col-0
Col-0, 27°C, 2 h
Col-0, 27°C, 24 h
Col-0, 37°C, 2 h
Col-0, 37°C, 24 h
cad2
cad2, 27°C, 2 h
cad2, 27°C, 24 h
cad2, 37°C, 2 h
cad2, 37°C, 24 h

Figure S6

A

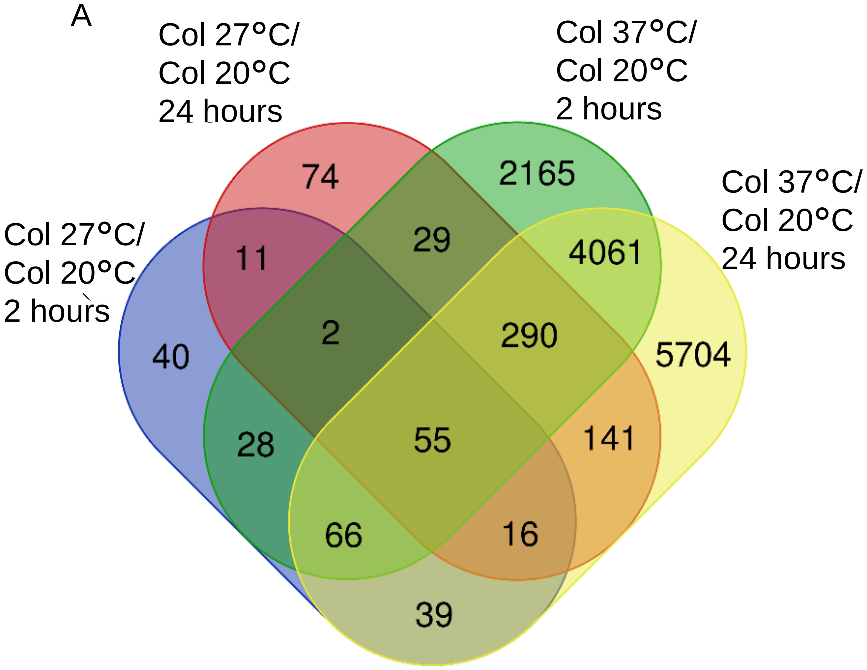


B

| GO category | p-value |
|---|---|
| 27°C specific | |
| Response to chemical stimulus | 5.1e-5 |
| 37°C specific | |
| Post embryonic development | 1.7e-21 |
| Response to stimulus | 3.7e-17 |
| Response to stress | 1.1e-9 |
| Response to chemical stimulus | 1.9e-7 |
| Post-translational protein modification | 4.7e-5 |
| Common | |
| Response to stimulus | 8.4e-21 |
| Response to chemical stimulus | 1.2e-18 |
| Response to stress | 6.0e-15 |
| Response to heat | 2.5e-11 |
| Response to temperature stimulus | 5.0e-11 |

C

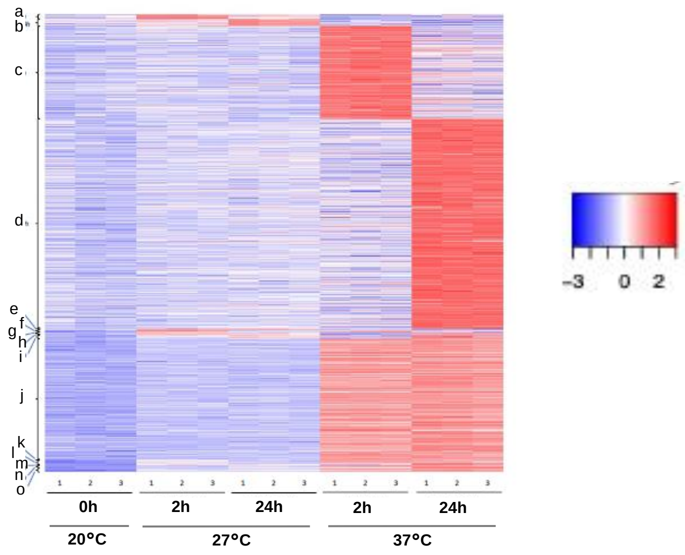


D

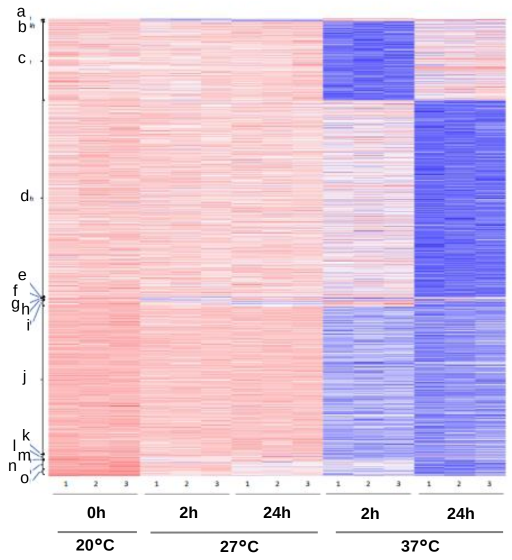

Figure S7

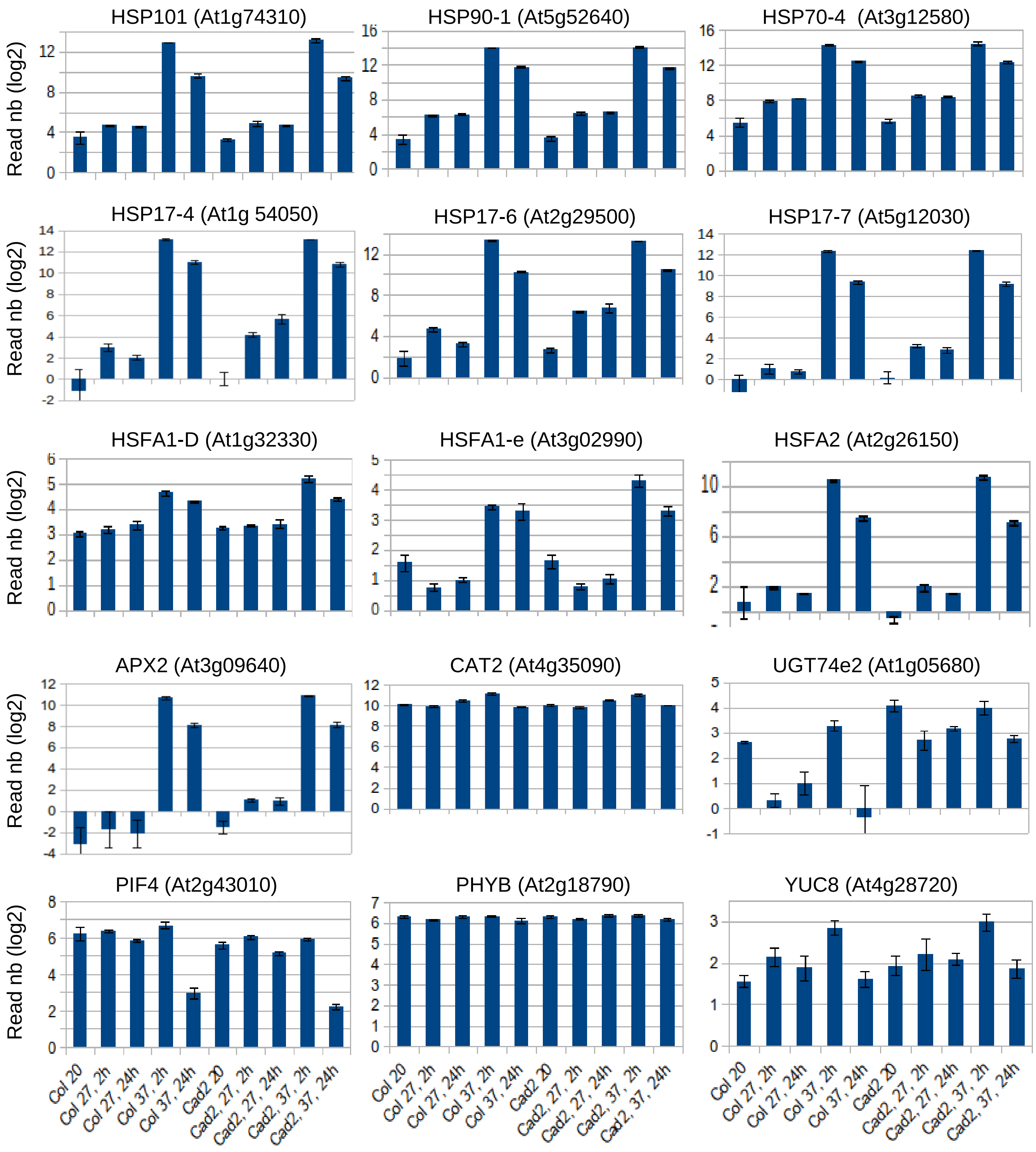
HSP101 (At1g74310)
HSP90-1 (At5g52640)
HSP70-4 (At3g12580)
HSP17-4 (At1g 54050)
HSP17-6 (At2g29500)
HSP17-7 (At5g12030)
HSFA1-D (At1g32330)
HSFA1-e (At3g02990)
HSFA2 (At2g26150)
APX2 (At3g09640)
CAT2 (At4g35090)
UGT74e2 (At1g05680)
PIF4 (At2g43010)
PHYB (At2g18790)
YUC8 (At4g28720)
Read nb (log2)
Col 20
Col 27, 2h
Col 27, 24h
Col 37, 2h
Col 37, 24h
Cad2 20
Cad2, 27, 2h
Cad2, 27, 24h
Cad2, 37, 2h
Cad2, 37, 24h

Figure S8

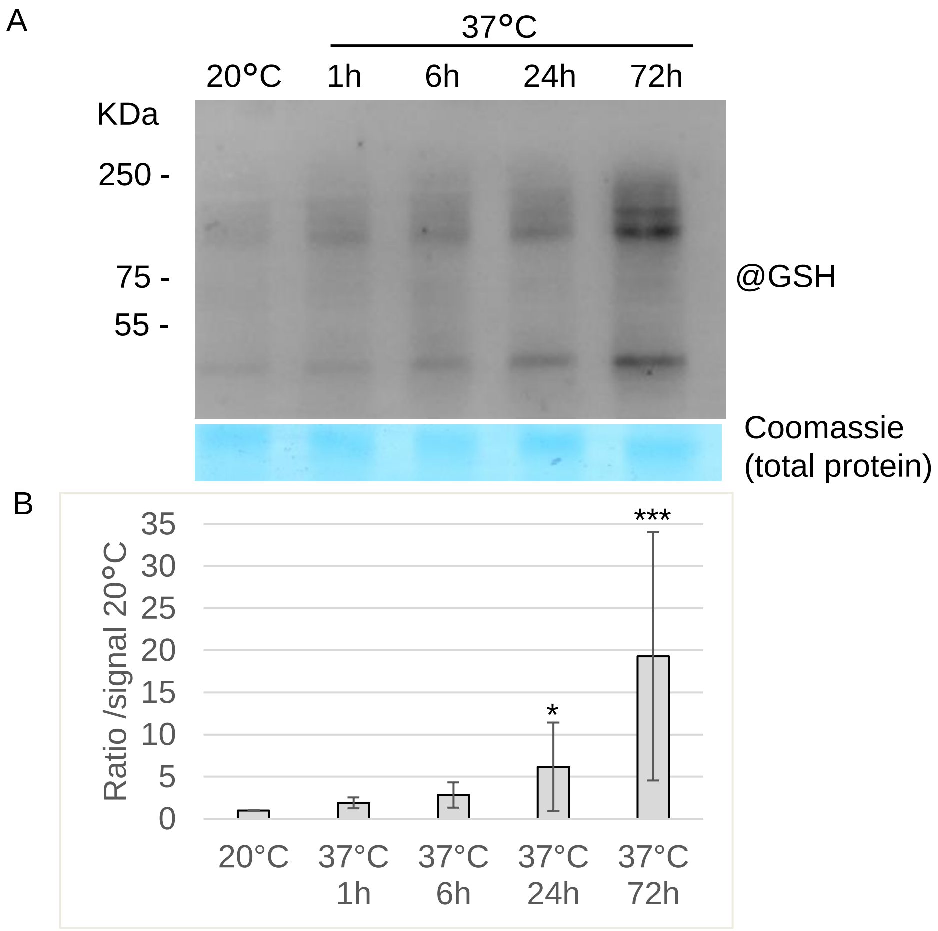

A
37°C
20°C
1h
6h
24h
72h
KDa
250 -
75 -
55 -
@GSH
Coomassie
(total protein)
B
Ratio /signal 20°C
35
30
25
20
15
10
5
0
***
*
20°C
37°C 1h
37°C 6h
37°C 24h
37°C 72h
